# B cell-specific XIST complex enforces X-inactivation and restrains atypical B cells

**DOI:** 10.1101/2021.01.03.425167

**Authors:** Bingfei Yu, Yanyan Qi, Rui Li, Quanming Shi, Ansuman Satpathy, Howard Y. Chang

**Affiliations:** Center for Personal Dynamic Regulomes, Stanford University, Stanford, CA, 94305, USA; Howard Hughes Medical Institute, Stanford University, Stanford, CA, 94305, USA; Department of Pathology, Stanford University, Stanford, CA, 94305, USA

## Abstract

The long noncoding RNA (lncRNA) XIST establishes X chromosome inactivation (XCI) in female cells in early development and thereafter is thought to be largely dispensable. Here we show XIST is continually required in adult human B cells to silence a subset of X-linked immune genes such as *TLR7*. XIST-dependent genes lack promoter DNA methylation and require continual XIST-dependent histone deacetylation. XIST RNA-directed proteomics and CRISPRi screen reveal distinctive somatic cell-specific XIST complexes, and identify TRIM28 that mediates Pol II pausing at promoters of X-linked genes in B cells. XIST dysregylation, reflected by escape of XIST-dependent genes, occurs in CD11c+ atypical memory B cells across single-cell transcriptome data in patients with female-biased autoimmunity and COVID-19 infection. XIST inactivation with TLR7 agonism suffices to promote isotype-switched atypical B cells. These results suggest cell-type-specific diversification of lncRNA-protein complexes increase lncRNA functionalities, and expand roles for XIST in sex-differences in biology and medicine.

**HIGHLIGHTS:**

- XIST prevents escape of genes with DNA hypomethylated promoters in B cells.
- XIST maintains X-inactivation through continuous deacetylation of H3K27ac.
- XIST ChIRP-MS and allelic CRISPRi screen reveal a B cell-specific XIST cofactor TRIM28.
- XIST loss and TLR7 stimulation promotes CD11c+ atypical B cell formation.

## INTRODUCTION

The inheritance of different X chromosome copies in XX females and XY males creates a challenge to equalize gene expression between the sexes in every cell of the body. Eutherian mammals have evolved a unique strategy to epigenetically silence one of the two X chromosomes in females, a process termed X chromosome inactivation (XCI) (Lyon, 1962). XCI is comprised of initiation, establishment and maintenance. During early embryogenesis, XCI is initiated by the expression of XIST, a long noncoding RNA (lncRNA) that is transcribed and spreads in *cis* across the entire inactive X chromosome (Xi) (Penny et al., 1996). XIST is essential for XCI establishment through a step-wise recruitment of multiple factors for transcriptional repression of the Xi (Chu et al., 2015; reviewed in Plath et al., 2002). Once XCI is established, the Xi is stably silenced through subsequent cell divisions via epigenetic mechanisms such as DNA methylation (Gartler and Riggs, 1983). Although most of X-linked genes are subjected to XCI, over 20% of X-linked genes can escape from XCI (termed escapee genes) in one or more cell types (reviewed in Carrel and Brown, 2017). Tissue-specific escapee genes have been recently discovered through genotype tissue expression project (GTEx), highlighting a diversity of XCI maintenance across different tissues (Oliva et al., 2020; Tukiainen et al., 2017). Despite a central role of XIST in the initiation and establishment process of XCI, XIST was long thought to be dispensable for XCI maintenance. Genetic deletion of XIST in cultured human or mouse somatic cells did not reactivate the Xi, based on a handful of transgenic or endogenous X-linked reporter genes (Brown and Willard, 1994; Csankovszki et al., 1999). However, recent studies with conditional inactivation of mouse *Xist* in different tissues showed partial Xi reactivation and increased escapee gene expression, with variable consequences for tissue function and organismal fitness (Adrianse et al., 2018; Yang et al., 2016, 2020). For instance, deletion of *Xist* in mouse hematopoietic stem cells invariably led to a myelodysplatic syndrome and ultimately leukemia (Yildirim et al., 2013), indicating the blood compartment is sensitive to Xist loss. These observations raise essential questions regarding the role of XIST in human XCI maintenance, and highlight the need to understand fundamental principles that can predict when and where XIST is needed for XCI maintenance.

XIST lncRNA (19 kb long in humans) comprises several functional modules including A-F repeats that are conserved between human and mouse (reviewed in Lu et al., 2017). These functional repeats in XIST RNA can bind and recruit ∼80 proteins, through both direct RNA-protein and indirect protein-protein interactions, in embryonic stem cells (Chu et al., 2015). The XIST associated proteins are responsible for chromatin modification, transcriptional silencing, XIST RNA spreading and coating, and chromosome-wide architecture reorganization (reviewed in Loda and Heard, 2019). At the onset of XCI, XIST is upregulated on the prospective Xi and recruits SPEN via A-repeat to facilitate histone deacetylation as early silencing events (Chu et al., 2015; Lu et al., 2016; Żylicz et al., 2019). Then XIST B and C repeat binding protein HNRNPK functions to recruit Polycomb repressive complex (PRC) 1 and 2, which are responsible for later accumulation of repressive histone marks H2AK119b and H3K27me3 respectively to further lock in silencing (reviewed in Almeida et al., 2020). XIST RNA tethering to the Xi relies on C repeat binding protein HNRNPU and E repeat binding protein CIZ1 since deletion of these proteins lead to a dispersed localization of XIST (Hasegawa et al., 2010; Sunwoo et al., 2017). XIST also plays a central role in reorganization of the 3D chromatin structure of the Xi which is devoid of topologically-associated domain and encompass a unique bipartite structure (Giorgetti et al., 2016; Rao et al., 2014). Additional XIST interacting proteins include RBM15 and WTAP that are critical for m6A methylation of XIST RNA, and LBR which is essential for the Xi anchoring to the nuclear lamina (Chen et al., 2016; Patil et al., 2016). Although XIST binding proteins are well characterized in the context of XCI establishment, the nature and function of the XIST ribonucleoprotein (RNP) complex during XCI maintenance in adult somatic cells is virtually unknown.

Escape from XCI has long been thought to contribute to sex differences in biology and medicine. Because the X chromosome harbors a high density of immune genes, incomplete XCI maintenance may lead to increased expression of escapee genes and elevated immune gene dosage in females over males, contributing to female-biased immune features such as higher B cell number and antibody production compared to male. Such heightened humoral responses are beneficial to fight against pathogen infections while can be detrimental via production of self-attacking autoantibodies, a hallmark of autoimmune disease (reviewed in Klein and Flanagan, 2016). Indeed, 80% of patients with autoimmune diseases are women (reviewed in Libert et al., 2010). Males with an extra X (Klinefelter syndrome, XXY) have 14 fold higher risk to develop systemic lupus erythematosus (SLE) than males with single X (XY), and females with a single X (Turner syndrome, XO) are not enriched in SLE patients (Cooney et al., 2009; Scofield et al., 2008). Thus, the female-biased autoimmunity is not solely due to gonadal sex or hormone differences, but is intimately linked to X chromosome dosage, especially the dosage of X-linked immune genes. For example, *TLR7* is an X-linked gene encoding Toll-like receptor 7, an innate immunity receptor that recognizes single-strand (ss)RNA-containing immune complexes, which are prevalent in female-biased autoimmunity and ssRNA viral infection (Celhar et al., 2012). The duplication of *Tlr7* gene in mice suffices to drive lupus-like symptoms whereas the deletion of *Tlr7* ameliorates the disease (Christensen et al., 2006; Subramanian et al., 2006).

An important consequence of elevated TLR7 signaling is the accumulation of CD11c+ atypical memory B cells, a unique B cell population recently recognized to greatly expand in aging, certain infectious diseases (malaria, HIV, and COVID19 (Woodruff et al., 2020)), and female-biased autoimmunity (SLE, rheumatoid arthritis (RA), and Sjögren’s syndrome (SS)) (reviewed in Cancro, 2020; Karnell et al., 2017). CD11c+ atypical B cells, also named age/autoimmune-associated B cells (ABC), double negative B cells, and tissue-like memory B cells, resides in a large pool of antigen-experienced B cells and differs from conventional memory B cells in phenotypic markers, activation requirements, differentiation pathway, and transcriptional programs (reviewed in Cancro, 2020; Karnell et al., 2017; Rubtsova et al., 2015). Atypical B cells specifically express T-bet, a master transcription factor for Th1 cytokines (Szabo et al., 2000), have skewed antibody isotype switching to IgG (Barnett et al., 2016; Wang et al., 2012), and their formation and activation depend on TLR7/9 signaling and Th1 cytokines, in contrast to conventional B cells that depend on BCR ligation and CD40 co-stimulation (Hao et al., 2011; Rubtsov et al., 2011; Rubtsova et al., 2015). Atypical B cells produce protective virus-specific antibodies in viral infections and pathogenic autoantibodies in the context of autoimmunity. (Barnett et al., 2016; reviewed in Cancro, 2020). Importantly, CD11c+ atypical B cells exhibit a strong female bias and selectively accumulate in aged female mice but not in age-matched male mice (Rubtsov et al., 2011). How and why this CD11c+ atypical B cell subset is female-biased and potential links to X chromosome dosage and XIST action are largely unknown. A recent study showed a complete lack of XIST localization on the Xi in CD11c+ atypical B cells compared to conventional memory B cells, suggesting a defect of XCI maintenance may occur in this unique B cell subset (Pyfrom et al., 2020). Such abnormal XIST localization has also been observed in patients with female-biased SLE where CD11c+ atypical B cells are massively accumulated (Pyfrom et al., 2020; Wang et al., 2016). The connection between female-bias, abnormal XIST localization, and hyper-responsiveness to TLR7 which is encoded by an X-linked immune gene in CD11+ atypical B cells, raised a possibility that the fidelity of XCI maintenance may be perturbed in these cells. Although correlative in nature, these observations appear contrary to the dogma that XIST is dispensable for XCI maintenance in somatic cells.

Here, we investigate the requirement and mechanism of action of XIST for XCI maintenance in human B cells. We identify X-linked genes that require ongoing XIST-mediated silencing, and use these gene loci to characterize specific epigenomic features *in cis* and XIST-associated factors *in trans* that orchestrate their silencing. We further track XIST dysregulation at single-cell level across multiple human disease states and link XIST dysregulation to CD11c+ atypical B cells. Gene editing in primary human B cells validate a role for XIST in restraining the formation of CD11c+ atypical B cells. These multiple lines of evidence show that XIST assembles adult cell-type specific complexes to achieve cell type-specific functions, suggesting a general mechanism to broaden and diversify lncRNA-mediated gene regulation.

## RESULTS

### XIST is essential for maintaining X-inactivation of immune genes in human B cells

We chose to work with GM12878 because it is an immortalized female human B cell line with normal karyotype, has been extensively characterized in the 1000 Genomes and ENCODE projects with a wealth of genomic and epigenomic data (1000 Genomes Project Consortium et al., 2012; ENCODE Project Consortium, 2012), and has a well-behaved and prototypical Xi structure representative of human and primate somatic cells (Giorgetti et al., 2016; Rao et al., 2014). To determine if the X-inactivation is maintained by XIST RNA-mediated silencing or epigenetic memory such as DNA methylation and H3K27me3, we performed CRISPR interference (CRISPRi) of XIST in dCas9-KRAB expressing GM12878 B cells (sgXIST group), leading to 75% knockdown (**Figure 1A-B, Figure S1A**). We also interfered with the epigenetic memory by treating the cells with DNMT and EZH2 inhibitors to simultaneously block DNA methylation and H3K27me3 (inhibitors group). To systemically investigate the XCI maintenance, we performed allelic RNA-seq analysis using informative single nucleotide polymorphisms (SNPs) provided by the fully phased genome of this B cell line (ENCODE Project Consortium, 2012). For genes with at least 10 reads that can be assigned to alleles, we calculated the allelic bias using d-score (reads ^Xi^/(reads^Xi^+ reads^Xa^)- 0.5). The d-score has a range of −0.5 to +0.5: -0.5 means all the reads are from the active X chromosome (Xa) and +0.5 means all the reads are from the inactive X (Xi); 0 means the reads are equally distributed between Xi and Xa alleles. Xa serves as an internal control to minimize the impact of experimental variance.

**Figure 1.**
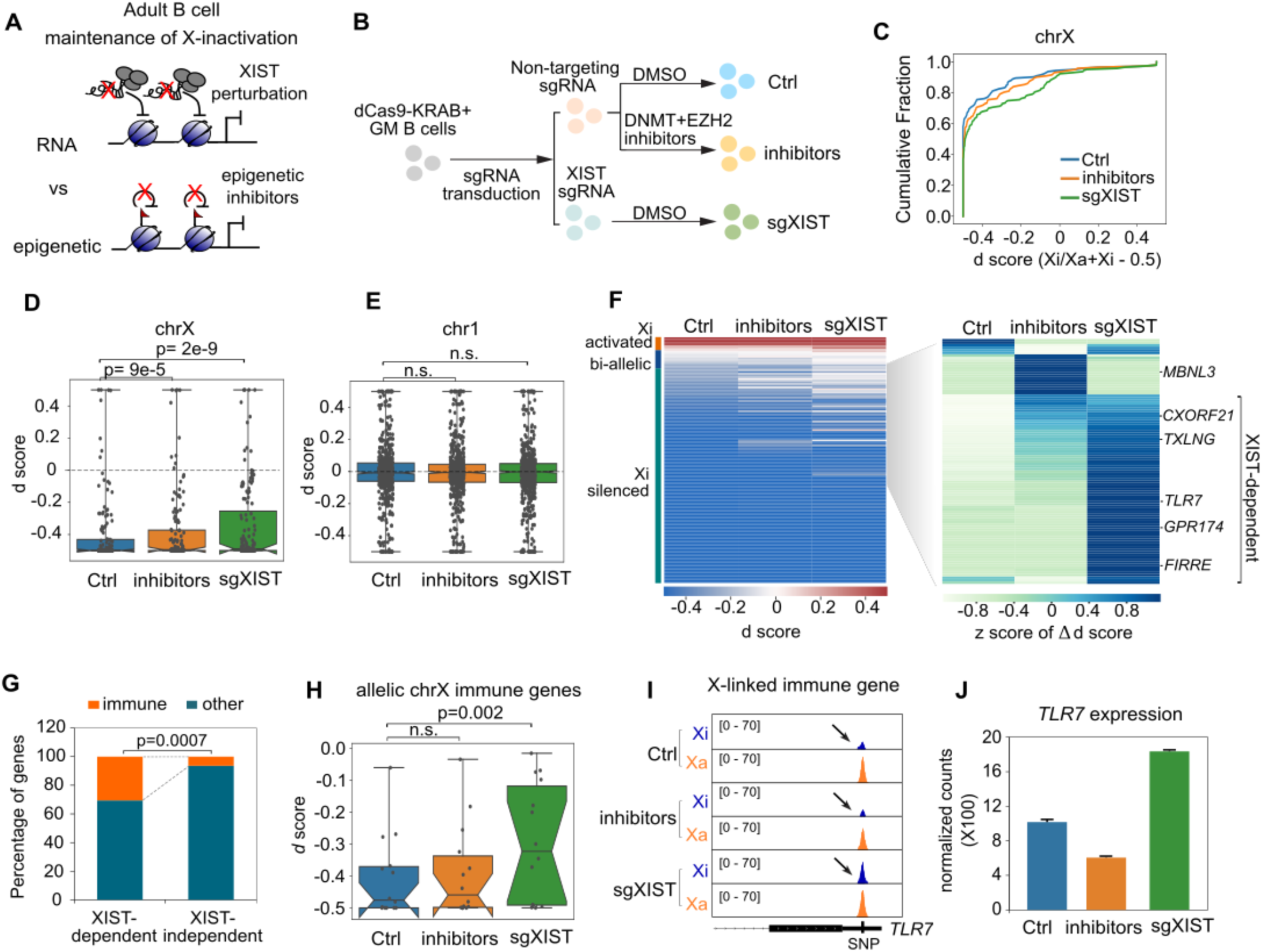
Xist is essential for XCI maintenance of a subset of X-linked genes. A. Schematic view of hypothesis. To test if XCI maintenance is controlled in Xist RNA-mediated mechanism or epigenetic memory mechanism, X chromosome (chrX) allelic expression were measured in B cells with XIST perturbation by CRISPRi or in B cells treated with DNA methylation and H3K27me3 inhibitors for 7 days. B. Schematic view of experimental set up. Control group: B cells perturbed with non-targeting control sgRNA and treated with DMSO for 7 days. Inhibitors group: B cells perturbed with non-targeting control sgRNA and treated with DNA methylation (5’-azacytidine) and Ezh2 (EPZ-6438) inhibitors for 7 days. sgXIST group: B cells perturbed with XIST sgRNA and treated with DMSO for 7 days. C. Cumulative distribution of d score of allelic gene expression across the X chromosome in ctrl, inhibitors and sgXIST group. P value was calculated using Kolmogorov–Smirnov test. D. Box plot showing the distribution of d score of allelic gene expression on chrX between control, inhibitors and sgXIST group. P values were calculated using paired t-test. E. Box plot showing the distribution of d score of allelic gene expression on chr1 between control, inhibitors and sgXIST group. P values were calculated using paired t-test. F. Left heatmap showing the d score of allelic gene expression on chrX between control, inhibitors and sgXIST group. Right heatmap showing the hierarchical clustering of genes with differential allelic expression between groups and reveals XIST-dependent genes whose d score are higher in sgXIST group compared to the control group. Value depicts the z score of the difference of d score. Representative XIST-dependent and -independent genes are highlighted. G. Bar plot showing the percentage of immune-related gene and genes with other functions between XIST-dependent and -independent genes. P value was calculated using Fisher exact test. H. Box plot showing the distribution of d score of allelic gene expression on X-linked immune genes between control, inhibitors and sgXIST group. P values were calculated using paired t-test. I. Allelic specific RNA-seq tracks showing the allelic expression of X-linked immune gene *TLR7*. Xa, active X chromosome. Xi, inactive X chromosome. J. Bar plot showing the RNA-seq normalized counts of *TLR7* expression between control, inhibitors and sgXIST groups.

Treatment of epigenetic inhibitors led to global change of transcriptional programs, ∼8000 differentially expressed genes (DEG) compared to control group. In contrast, silencing of XIST leads to only ∼500 DEGs (**Figure S1C, Tabel S1**). Despite the difference in DEG numbers, we noticed that upregulated DEGs after XIST knockdown are more enriched on X chromosome (24.4%) compared to inhibitors-treated cells (6.8%) (**Figure S1D**). These results indicated that XIST is the major contributor to maintain the gene dosage on X in somatic B cells, which is also supported by the Pearson correlation coefficient analysis of the allelic bias on the X across all samples (**Figure S1B**). The cumulative distribution of the d score across the X showed a significantly increased d score in sgXIST group compared to control group (*p*=0.002, K-S test). Further allelic analysis showed that reduction in XIST leads to significant Xi reactivation of X-linked genes whereas there are no changes of allelic bias on autosomes such as chromosome 1 (**Figure 1D-E, Figure S1E**). For 130 genes with lower expression in Xi (d-score <0), a subset of genes (27.6%, including *TLR7, FIRRE, CXORF21, TXLNG*) showed significantly increased d-score in sgXIST cells compared to control cells (d score ^sgXIST^- d score^Ctrl^ >0.03, binomial *p*<0.05), suggesting that they rely on XIST to maintain X-inactivation (**Figure 1F**). Thus, contrary to expectation, XIST is not fully dispensable for XCI maintenance in adult B cells.

We named this subset of 36 genes as XIST-dependent genes, whereas additional 94 X-linked genes whose d-score are not significantly altered after XIST perturbation were named as XIST-independent genes (**Table S2**). In control B cells, the d-score is significantly higher in XIST-dependent compared to -independent genes, suggesting that these XIST-dependent genes are not stably silenced (**Figure S1F**). Indeed, we compared the proportion of variable and constitutive escapee genes that are defined from GTEx study (Tukiainen et al., 2017) and found these escapee genes are enriched in XIST-dependent compared to -independent genes (**Figure S1G**). Gene Ontology analysis of genes that are upregulated in sgXIST group showed enrichment of immune response including immune genes like *TLR7, GPR174, CXCR3, CR2, CXCL10, IFIT3, and OAS1* (GO enrichment *p*=7 x 10^-5^) (**Figure S1H**). Immune-associated genes are also more enriched in XIST-dependent genes compared to -independent genes (*p*=0.01, Fisher’s exact test) (**Figure 1G**). These results suggested that XIST may be essential for balancing the gene dosage of X-linked immune genes. As expected, we found increased d-score of X-linked immune genes in sgXIST group compared to control group, supporting the idea that XIST is essential for Xi silencing of these immune-associated X-linked genes (**Figure 1H**). For example, *TLR7* is an X-linked gene encoding Toll-like receptor 7, a key innate immunity receptor for single-strand RNA, whose gene dosage and biallelic expression are associated with female-biased autoimmune diseases (Deane et al., 2007; Souyris et al., 2018). We observed an increased biallelic expression of *TLR7* and ∼1.8 fold increase in *TLR7* bulk RNA expression after XIST knockdown (**Figure 1I-J**).

To further test if XIST is able to re-silence active escapee genes in B cells, we restored XIST expression in XIST CRISPRi B cells via introduction of an anti-CRISPR protein. qRT-PCR showed ∼90% restoration of XIST expression in sgXIST B cells (**Figure S1I**). Allelic RNA-seq analysis showed significantly decreased d scores across the X chromosome in sgXIST+anti-CRISPR cells compared to sgXIST alone cells (**Figure S1J-K**), indicating the restoration of XIST expression potently re-silenced previously active escapee genes in adult somatic B cells.

Together, we showed that XIST is essential and continuously required to maintain the Xi silence of a set of immune-associated X-linked genes in B cells.

### XIST requirement in XCI maintenance in other somatic cells

To understand the generality of XIST for XCI maintenance in other adult somatic cell types, we explored the XIST requirement in human female adult fibroblast cells (XIST CRISPRi in IMR90 cell line) and mouse female adult neuronal cells (re-analyzed RNA-seq of brain tissue from brain-specific XIST KO mice, (Carrette et al., 2018)). With comparable level of reduction of XIST expression in B cells, fibroblasts, and neuronal cells (**Figure S2A**), we found that XIST inactivation lead to upregulation of gene expression on the X compared to the autosome in all three cell lineages albeit with different magnitude (**Figure S2B**). XIST-silenced genes (the X-linked genes that are significantly upregulated after the loss of XIST) comprise of 14 genes in Ihuman fibroblasts and 19 genes in mouse neuronal cells, compared to 118 genes in human B cells. Among the XIST-silenced X-linked genes in fibroblasts and neuronal cells, several genes are variable escapees with tissue specificity revealed by GTEx analysis (**Figure S2 C-D**). Although there are only a modest number of genes that require XIST for XCI maintenance in fibroblasts and neuronal cells, different genes escape XCI in a tissue-specific manner and hence the number of potential XCI escapees increases with more tissues examined. For example, *Col4a5*, a type IV collagen gene whose mutation is associated with abnormal axogenesis in zebrafish (Takeuchi et al., 2015), is an Xist-silenced gene in mouse brain but not in fibroblasts or B cells (**Figure S2C**). Interestingly, *Col4a5* is also a brain tissue-specific escapee shown in human GTEx analysis (**Figure S2D**). The connection between cell type-specific XIST-silenced genes and tissue-specific escapees suggest that XIST may coordinate with cell/tissue-specific epigenetic environment to ensure XCI maintenance. Collectively, our data indicate that ongoing requirement for XIST to prevent escape from X inactivation is a general phenomenon, but B cells are most dependent on XIST with the largest number of genes among the three cell types we analyzed.

### Absence of DNA methylation predicts requirement of XIST for XCI maintenance

We hypothesized that XIST-dependent genes may harbor unique chromatin and/or genetic features associated with their unstable silencing and thus continuous dependence on XIST-mediated transcriptional repression. We first compared histone modifications, chromatin accessibility, and DNA methylation between XIST-dependent and -independent genes. We found significantly lower DNA methylation at TSS adjacent regions, and lower H3K27me3 deposition at the gene body of the XIST-dependent genes including *TLR7*, compared to XIST-independent genes such as *MBNL3* (**Figure 2A-B**). These differences may be a consequence of different transcriptional statuses or genomic features. Analysis of additional genomic features showed that XIST-dependent genes contain significantly fewer CpG islands (**Figure S2E**). DNA GC content, proximity of target gene loci to *XIST* locus, density of LINE, SINE, or LTR element were not significantly different between XIST-dependent and -independent genes (**Figure S2E-F**).

**Figure 2.**
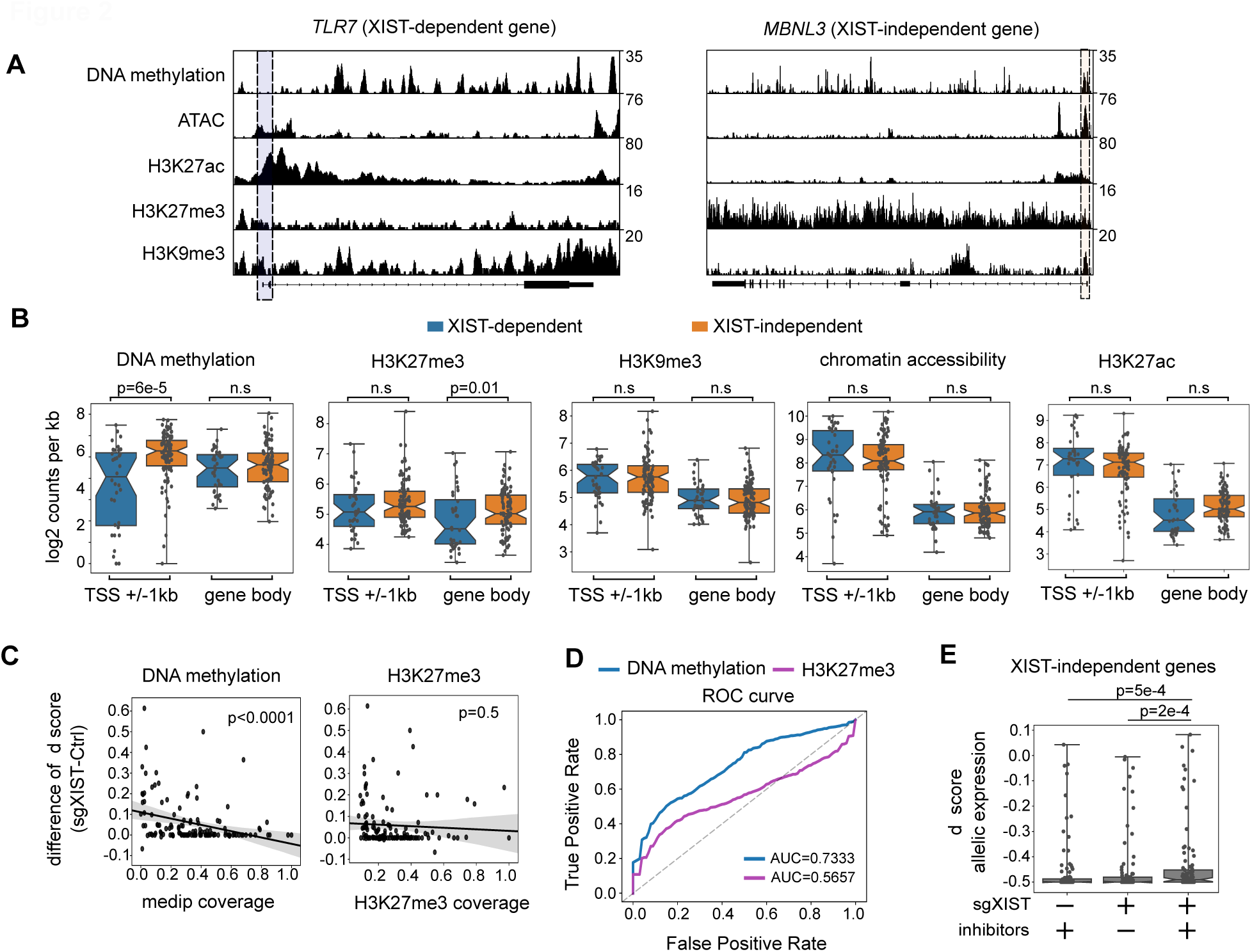
Low DNA methylation is associated with genes that continuously rely on Xist for XCI maintenance. A. Genome tracks showing chromatin accessibility (ATAC-seq), DNA methylation (medip-seq), ChIP-seq of H3K27ac, H3K27me3 and H3K9me3 profiles of representative XIST-dependent (*TLR7*) and -independent (*MBNL3*) genes. Shaded area highlights the promoter regions (TSS+/-1kb). B. Comparison of epigenetic features including DNA methylation (medip-seq), H3K27ac, H3K27me3, chromatin accessibility (ATAC-seq) and H3K9me3 between XIST-dependent and -independent genes. P value was calculated using non-parametric Mann-Whitney test. C. Scatter plots showing the linear regression analysis of relationship between XIST dependence (dscore^sgXIST^-dscore^ctrl^) and epigenetic features such as DNA methylation and H3K27me3 D. ROC curve of logistic regression analysis showing the prediction of XIST dependence using epigenetic features DNA methylation and H3K27me3. E. Box plot showing the d score of XIST-independent genes between inhibitors, sgXIST, and the combination of inhibitors treatment and XIST perturbation group. P values were calculated using paired t-test.

We further delineate the association among epigenomic features and XIST dependency using linear regression, and found that low DNA methylation is significantly associated with the genes whose XCI maintenance is dependent on XIST (**Figure 2C**). Notably, most of the XIST-dependent genes are in the lowest 10-percentile for promoter DNA methylation (**Figure 2C**). Logistic regression analysis indicated a key role of DNA methylation to predict whether the XCI maintenance of genes relies on XIST (**Figure 2D**). This suggests that the paucity of chromatin-based marks of epigenetic memory, specifically DNA methylation, increases the dependency on XIST for XCI maintenance. To determine the redundancy between epigenetic memory and XIST requirement for XCI maintenance, we treated XIST-depleted cells with epigenetic inhibitors to block DNA methylation and H3K27me3 and test if XIST-independent genes may become dependent on XIST for XCI maintenance when epigenetic memory is interfered. Indeed, we found significantly increased d score of XIST-independent genes in XIST depleted cells that are treated with DNMT and EZH2 inhibitors compared to either sgXIST group or inhibitors treated group (**Figure 2E**). Together, we show that XIST-dependent genes are not stably silenced due to the absence of DNA methylation. Moreover, XIST is functionally redundant with DNA methylation for the maintenance of silencing for larger set of Xi genes.

### XCI maintenance in adult B cells relies on XIST-mediated histone deacetylation

Next, we set out to investigate how XIST regulates XCI maintenance. XIST is critical for XCI initiation by recruiting multiple chromatin modification enzymes to the Xi, leading to the deacetylation of active mark H3K27ac and the accumulation of repressive marks including H3K27me3. To test if the loss of XIST impacts H3K27ac and H3K27me3 occupancy at Xi during XCI maintenance, we performed ChIP-seq of H3K27ac and CUT&RUN of H3K27me3 in a *XIST* knockout human B cell line. Using CRISPR-Cas9, we generated *XIST* A-repeat KO cells as a *XIST* KO model since the genomic deletion of *XIST* A repeat leads to >99% loss of XIST RNA (**Figure S3A-B**). We noticed that A-repeat is overlapped with the active promoter of *XIST*, suggesting the deletion of A-repeat may interrupt the *XIST* promoter and thus lead to a dramatic decrease of *XIST* RNA expression, which is consistent with the Xist A repeat KO mouse study (Hoki et al., 2009) (**Figure S3B-C**). Analysis of H3K27ac and H3K27me3 profiles showed a significant increase of H3K27ac deposition at Xi when XIST is ablated, while autosomes such as chromosome 1 showed no change (**Figure 3B**, **Figure S3D**), For example, compared to the control group, *XIST* KO led to increased H3K27ac deposition at Xi of the X-linked immune gene *GPR174* which has been recently shown to impart sex differences in B cell migration and humoral immunity (Zhao et al., 2020) (**Figure 3A**). This re-acetylation of H3K27 after the loss of XIST is preferentially enriched at XIST-dependent genes, suggesting that XCI maintenance of XIST-dependent genes is regulated by XIST-mediated H3K27ac deacetylation (**Figure 3B**). Notably, the re-acetylation of H3K27ac in *XIST* KO is significantly enriched at distal enhancers rather than promoter regions (**Figure 3C**). This is consistent with the recent discovery that during XCI initiation, XIST specifically promotes enhancer deacetylation by increasing the activity of histone deacetylase HDAC3, which pre-bound to those enhancers before XIST binding (Żylicz et al., 2019). Despite a loss of H3K27me3 at a few genes such as *FIRRE* in *XIST* KO cells (**Figure S3E**), we did not observe a significant difference of H3K27me3 accumulation at Xi between control and *XIST* KO cells, neither for its occupancy at XIST-dependent genes (**Figure 3D**). Thus, our data support the idea that XIST is continuously required to maintain the silencing status of the Xi especially for XIST-dependent genes in part through H3K27ac deacetylation at enhancers in B cells.

**Figure 3.**
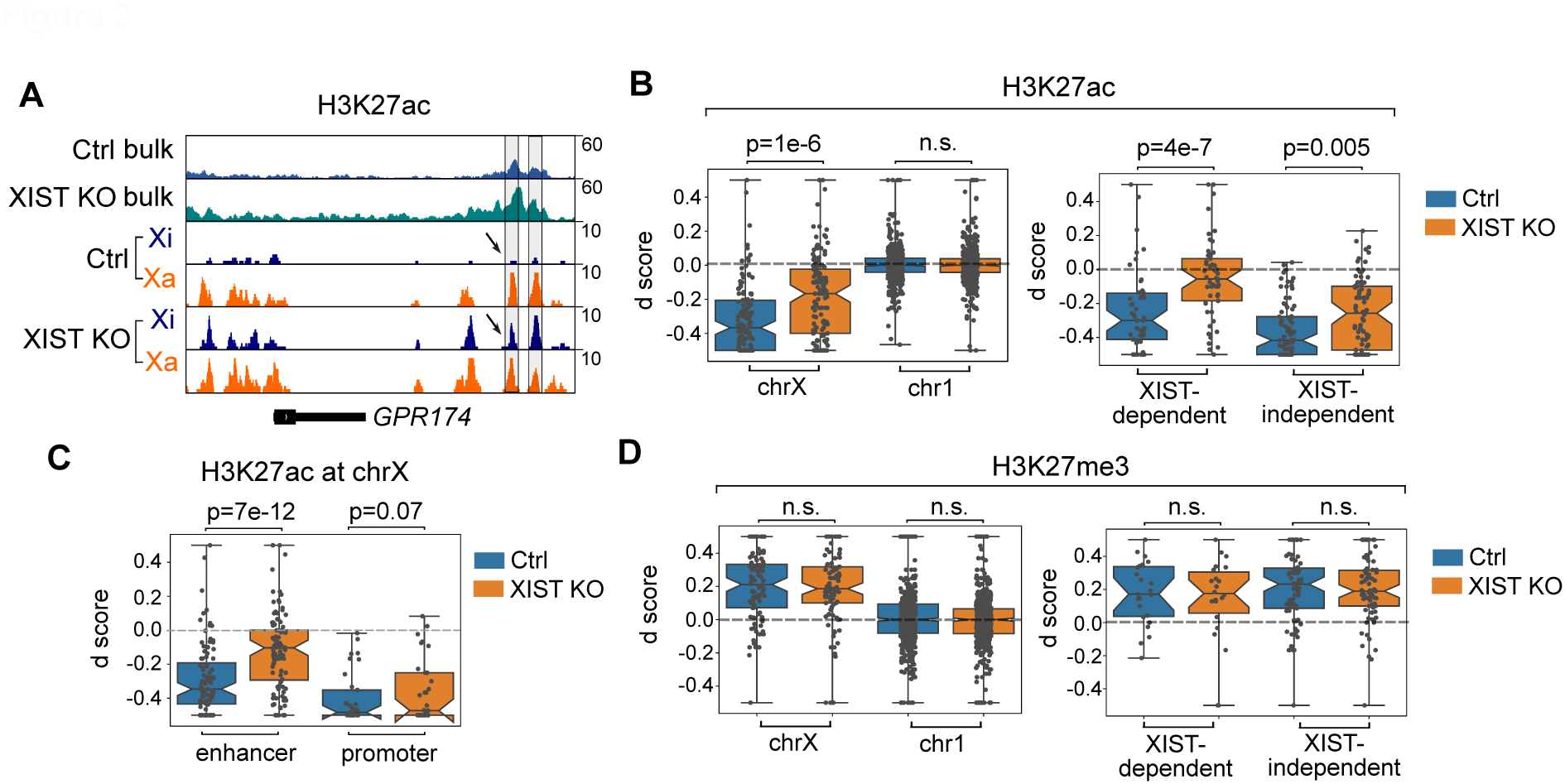
XIST mediates deacetylation of H3K27ac during XCI maintenance in somatic B cell line. A. H3K27ac ChIP-seq track of XIST-dependent gene *GPR174* between control and XIST KO group. B. Box plot (left) showing the d score of chrX and chr1 genes between H3K27ac allelic profiles of control and XIST KO group. Box plot (right) showing the d score of XIST-dependent and -independent genes between H3K27ac allelic profiles of control and XIST KO group. C. Box plot showing the H3K27ac allelic bias at enhancer and promoter regions of chrX genes between control and XIST KO cells. D. Box plot (left) showing the d score of chrX and chr1 genes between H3K27me3 allelic profiles of control and XIST KO group. Box plot (right) showing the d score of XIST-dependent and -independent genes between H3K27me3 allelic profiles of control and XIST KO group. All P values were calculated by paired t-test.

### XIST ChIRP-MS uncovers distinct somatic cell XIST RNP complexes for XCI maintenance

XIST is a modular lncRNA that can bind to diverse protein partners responsible for epigenetic silencing, nuclear matrix anchoring, and X chromosome spreading. These XIST bound co-factors involved in the XCI initiation have been recently identified by ChIRP-MS and RAP-MS (Chu et al., 2015; McHugh et al., 2015; Minajigi et al., 2015). However, the composition of the XIST ribonucleoprotein (RNP) complex after XCI establishment is not known in any adult somatic cells. To identify XIST co-factors, we performed XIST ChIRP-MS in the B cell line GM12878 and myeloid cell line K562, which both represent somatic cell types from the blood in which XCI has already been established (**Figure 4A, Table S3**). We used formaldehyde crosslinking that allows us to detect both direct RNA-protein interactions and indirect protein-protein interactions in the XIST RNP complex. We added an RNAse treatment group as a negative control to avoid the non-specific proteins that bind to probes and streptavidin beads. The XIST probes retrieved 60% of XIST RNA without contamination of abundant housekeeping gene *GAPDH* mRNA, indicative of a good specificity for these probes (**Figure S4A**). Additionally, XIST probes retrieved numerous proteins that are quite abundant as visualized by Coomassie Blue staining (**Figure S4B**).

**Figure 4.**
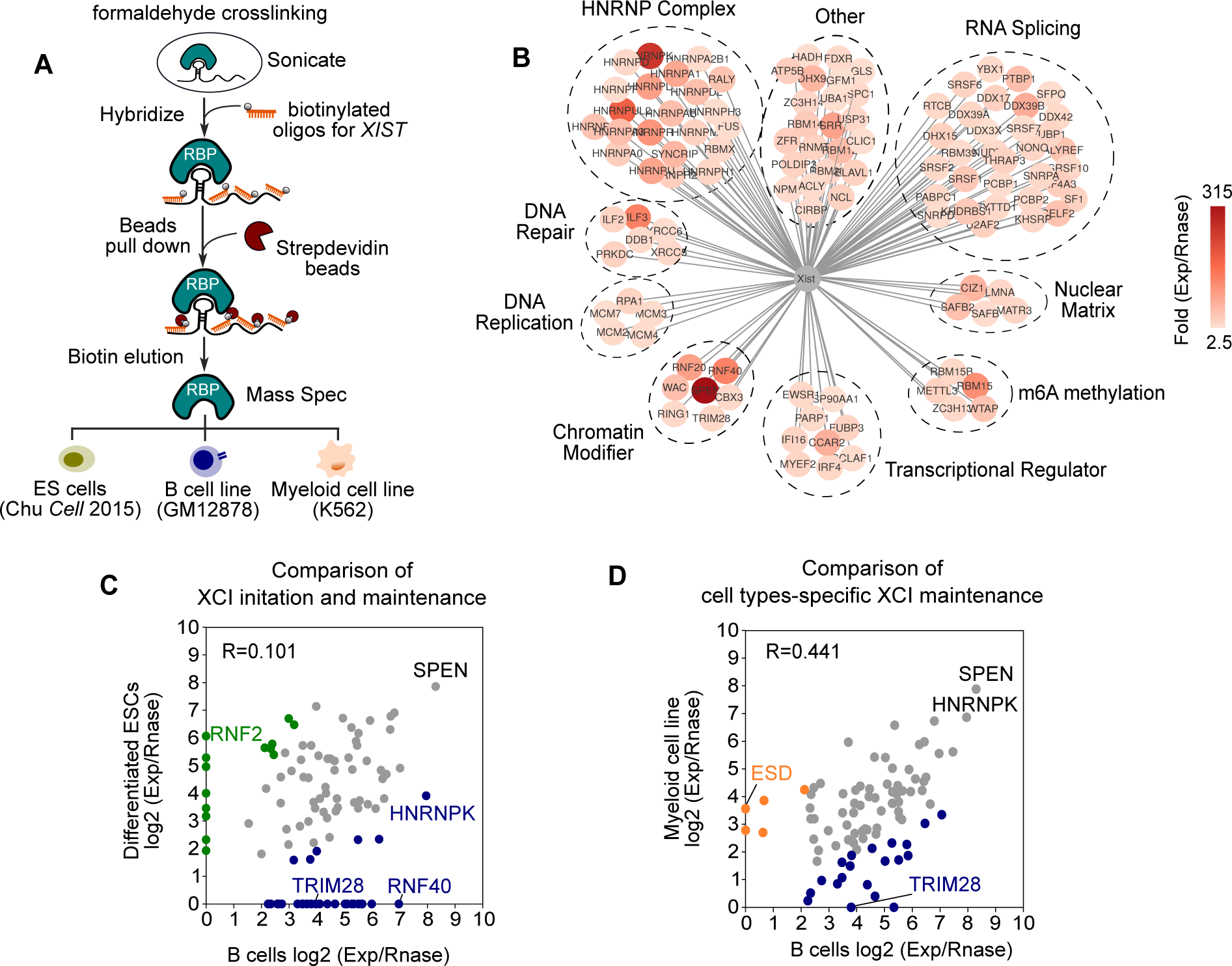
Identification of a unique XIST RNP complex in somatic B cell line by XIST CHIRP-MS. A. Schematic view of ChIRP-MS of XIST bound proteins in ES cell line, GM12878 B cell line, and K562 myeloid cell line. B. XIST RNA-associated protein network in B cell line revealed by Cytoscape. Proteins are with known functions or belonging to known complexes are grouped and annotated together. Color and value depicts the fold change of peptide counts of experimental group compared to Rnase treated group. C. Scatter plot showing the comparison of XIST-bound protein enrichment between differentiated ES cell line and B cell line. B cell enriched proteins (>2 fold in B cells versus ES cells) are labeled in blue; ES enriched proteins are labeled in green. D. Scatter plot showing the comparison of XIST-bound protein enrichment between myeloid cell line and B cell line. B cell enriched proteins (>2 fold in B cells versus myeloid cells) are labeled in blue; myeloid enriched proteins are labeled in orange.

Using two replicates of XIST ChIRP-MS data in B cell line, we found ∼100 XIST binding proteins that are enriched in experimental group but not in RNAse treatment group and are potentially important for XCI maintenance in B cells (**Figure S4C**). These XIST co-factors can be segregated into groups based on their functions. The groups include chromatin enzymes, nuclear matrix, m6A methylation, and RNA splicing, etc, which may serve diverse roles for X-inactivation (**Figure 4B**). For example, the transcriptional repressor SPEN is recruited to Xi by XIST A-repeat and promotes gene silencing via modulating HDAC activity (Chu et al., 2015; Dossin et al., 2020; McHugh et al., 2015). RBM15 has been shown to regulate m6A methylation of XIST to facilitate XIST-mediated transcriptional repression (Patil et al., 2016). In addition to the known XIST co-factors, we found novel XIST binding proteins in B cells such as TRIM28 (also known as KAP1), an epigenetic co-repressor that can bridge multiple histone modifiers like SETDB1 for epigenetic repression (Czerwińska et al., 2017; Iyengar and Farnham, 2011). To determine if the XIST-associated RNP complex is similar between XCI initiation and maintenance, we compared the enrichment of peptide counts for XIST binding proteins between B cells and ES cells with inducible expression of XIST. We found 57.8% of XIST-binding proteins are shared between XCI initiation and maintenance (**Figure 4C**). This comparison revealed a list of XCI maintenance-specific XIST RNP subunits including RNF40 and TRIM28, suggesting XIST may orchestrate a network of proteins specifically for XCI maintenance. We further compared XIST binding proteins between B cells and myeloid cells and found that 71.3% of XIST binding proteins are shared between these two somatic cell types, suggesting that XIST-RNP also has cell-type specific subunits in adult somatic cells. Notably, this comparison revealed that TRIM28 is a B cell-specific XIST binding co-factor, suggesting that TRIM28 might be essential for XIST-mediated XCI maintenance in B cells (**Figure 4D**). To further validate TRIM28 as a B cell-specific XIST cofactor, we performed TRIM28 RIP-qRT-PCR with formaldehyde crosslinking in B cells, K562 myleoid cells, and differentiated ES cells (**Figure S4D**). Despite similar TRIM28 immunoprecipitation efficiency among all three cell types, TRIM28 specifically retrieved XIST RNA only in B cells but not in myeloid cells or ESCs, indicating that TRIM28 is a B cell-specific XIST binding protein. Collectively, we mapped the XIST interactome in somatic immune cells and identified a catalog of XIST binding proteins that are potentially critical for XCI maintenance.

### Allelic CRISPRi screen reveals essential XIST co-factors for XCI maintenance in B cells

To functionally validate if the XIST binding proteins revealed by ChIRP-MS are essential for XCI maintenance, we developed a CRISPRi screen in the dCas9-KRAB expressed B cell line. The readout of this assay is the allelic expression of *TLR7*, an X-linked immune gene whose biallelic expression is associated with female-biased autoimmune disease. We first selected 57 high-confidence XIST-associated proteins in B cells with a stringent cutoff (peptide counts>25, fold enrichment >10) and designed two sgRNAs per target gene with each targeting a different region in 200 bp regions downstream TSS. To increase the CRISPRi efficiency, we cloned two sgRNAs under human or mouse U6 promoters in the same lentiviral vector. After individual sgRNA transduction and puromycin selection, we enriched for cells undergoing CRISPRi and isolated RNA for *TLR7* allelic library preparation (**Figure 5A, Table S4**).

**Figure 5.**
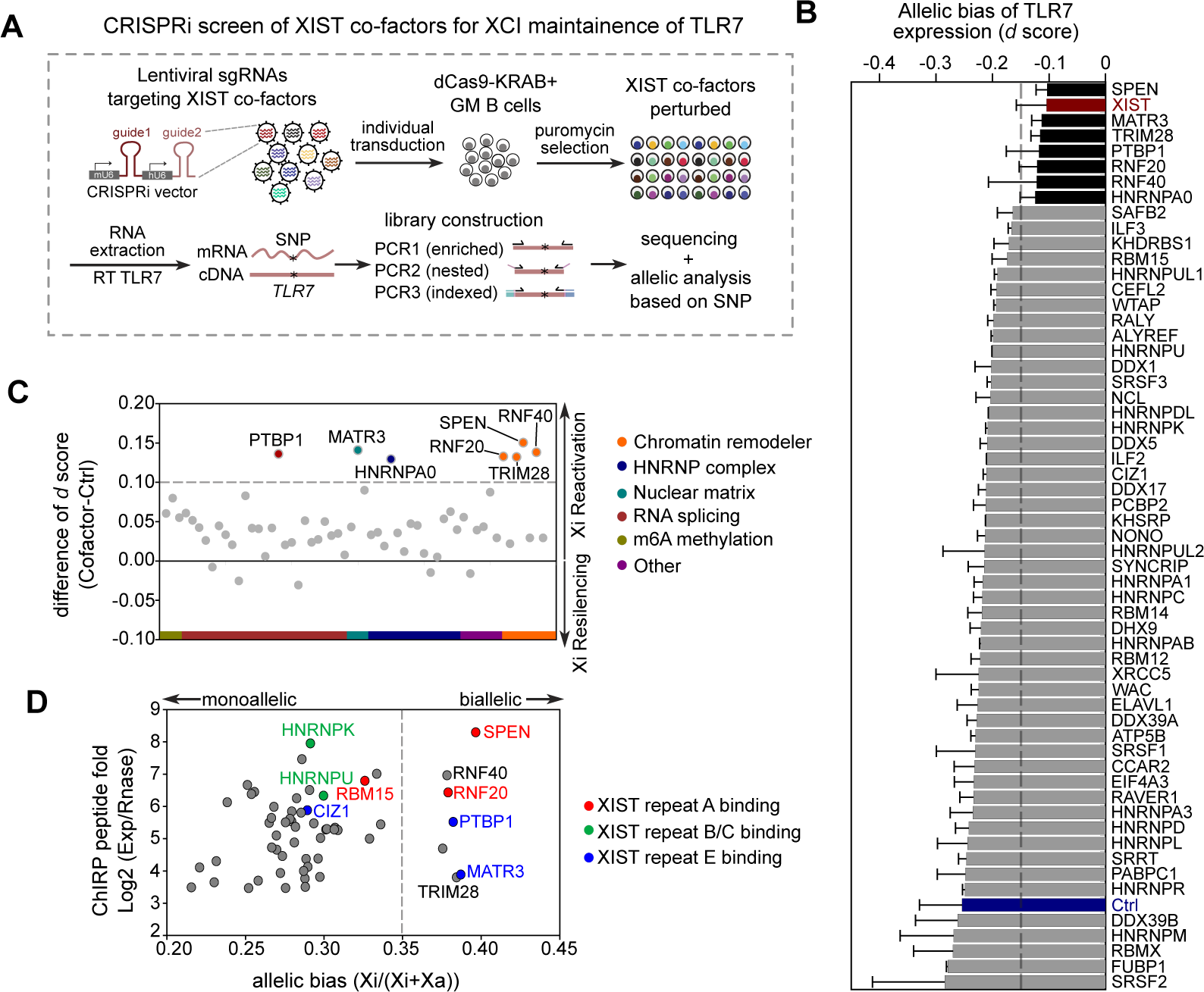
Allelic CRISPRi screen reveals key XIST cofactors responsible for XCI maintenance of X-linked immune gene *TLR7*. A. Schematic view of CRISPRi screen of highly-confident XIST cofactors that impact *TLR7* allelic gene expression. B. Bar plot showing the mean d score of TLR7 from two replicate screens after individual perturbation of XIST cofactors. Negative control (Ctrl) is labeled in blue and positive control (sgXIST) is labeled in red. C. Same as in B. The enriched co-factors compared to negative control are grouped in different functions or complexes and highlighted with different colors. D. Scatter plot showing the co-factors with their XIST binding signal shown by fold change of peptide counts and their effect on *TLR7* allelic gene expression shown by allelic bias score. Known XIST repeat A, B/C and E binding factors are highlighted in different colors. Gray dashed line in B-D indicates cut-off of the difference of d score: d score ^co-factor^-d score^ctrl^=0.1

Similar to *XIST*-depleted cells as a positive control, depletion of 7 out of 57 XIST cofactors individually showed an increased d-score of *TLR7* compared to the negative control group using non-targeting sgRNA, suggesting that these co-factors are critical for XCI maintenance of *TLR7* (cut-off delta d score: d score^co-factor^ - d score^Ctrl^ >0.1)(**Figure 5B**). The 7 key XIST co-factors are SPEN, TRIM28, RNF20, RNF40, PTBP1, MATR3, and HNRNPA0, and the first four 4 proteins are involved in chromatin modification (**Figure 5C**). For example, SPEN, a well-established XIST co-factor that facilitates histone deacetylation critical for XCI initiation, is also essential for XCI maintenance of *TLR7*. This is consistent with our discovery that XIST can modulate XCI maintenance through histone deacetylation, suggesting that XIST can recruit SPEN to promote histone deacetylation to maintain Xi status. Notably, despite similar XIST binding affinity and perturbation efficiency with XIST A-repeat binding factors like SPEN, depletion of XIST repeat B or C binding factors such as HNRNPK and HNRNPU did not affect XCI maintenance. (**Figure 5D, Figure S5A**). HNRNPK has been shown to recruit Polycomb repressive complexes for H2AK119ub and H3K27me3 spreading across Xi during XCI establishment (Chu et al., 2015; Colognori et al., 2019; Pintacuda et al., 2017). Given few change of H3K27me3 on Xi in the absence of XIST and the modest effect of HNRNPK on XCI maintenance (**Figure 3E, Figure 5D**), the results indicated that HNRNPK-recruited Polycomb complexes and subsequent H3K27me3 deposition may be dispensable for XCI maintenance, highlighting that XIST-recruited SPEN and SPEN-meidated H3K27ac deactylation might be the major factor to ensure the XCI maintenance. To further confirm the role of SPEN in XCI maintenance of B cells, we generated a SPEN mutant B cell line with deletion of SPEN RRM2-4 domain, which directly interacts with XIST to mediate gene silencing (Carter et al., 2020; Dossin et al., 2020; Monfort et al., 2015)(**Figure S5B**). Allelic analysis of SPEN mutant showed that the loss of function of SPEN led to significant reactivation of Xi specifically at XIST-dependent genes and largely recapitulated the effect of XIST inactivation (**Figure S5C-D**). These data fully support the idea that XIST recruits SPEN to maintain the XCI likely via SPEN-HDAC3 mediated deacetylation of H3K27ac.

PTBP1 is involved in XIST splicing (Stork et al., 2019), and MATR3 is a nuclear matrix protein potentially involved in XIST nuclear retention. Both PTBP1 and MATR3 have been recently reported to play a key role in XCI maintenance (Pandya-Jones et al., 2020). Importantly, PTBP1 and MATR3 are not required for initiation of XCI but only for its maintenance in ES cells (Pandya-Jones et al., 2020), which is fully consistent with our discovery of both factors as required to restrain escape from XCI in adult B cells. Importantly, depletion of B cell-specific XIST co-factor TRIM28 led to Xi reactivation of *TLR7*, indicating that potential TRIM28-mediated mechanisms participate in XCI maintenance (**Figure 5C**). Together, our screen uncovered XIST binding co-factors that are critical for XCI maintenance of *TLR7* in B cells, providing insights into a potential mechanism for XIST-mediated XCI maintenance.

### TRIM28 is required for B cell XCI maintenance of a subset of X-linked genes

Our ChIRP-MS data and allelic CRISPRi screen unveiled TRIM28, a novel B cell-specific XIST RNA-associated protein, that is essential for XCI maintenance of *TLR7*. To determine if TRIM28 is critical for XCI maintenance of other X-linked genes, we generated *TRIM28* KO B cells, which reduced TRIM28 protein by >95% (**Figure S6A**). Consistent with the screen data, we found Xi reactivation of *TLR7* in *TRIM28*-KO cells (**Figure 6A**). However, d-score analysis of the entire X chromosome showed a modest effect on Xi reactivation in *TRIM28* KO cells (**Figure 6B**). It is likely that *TRIM28* is responsible for XCI maintenance of a small subset of genes such as *TLR7*. Indeed, clustering analysis showed that the loss of TRIM28 leads to increased d score of a subset of genes and 64.7% of them are also XIST-dependent (**Figure 6C**). The increased d score of these TRIM28-dependent genes (d score^TRIM28 KO^- d score^Ctrl^ >0.03) is due to re-activation of Xi without significant difference on Xa (**Figure S6B**). To examine whether the TRIM28-dependent genes are directly occupied by TRIM28, we performed TRIM28 ChIP-seq in this B cell line. On autosomes, TRIM28 occupied the ZNF genes with high modification levels of the repressive mark H3K9me3 (**Figure S6C**), as expected based on prior studies (Iyengar and Farnham, 2011). On the X, TRIM28 is enriched at the promoters of TRIM28-dependent X-linked genes, such as *TXLNG* (**Figure 6D**), compared to TRIM28-independent genes (**Figure 6E**).

**Figure 6.**
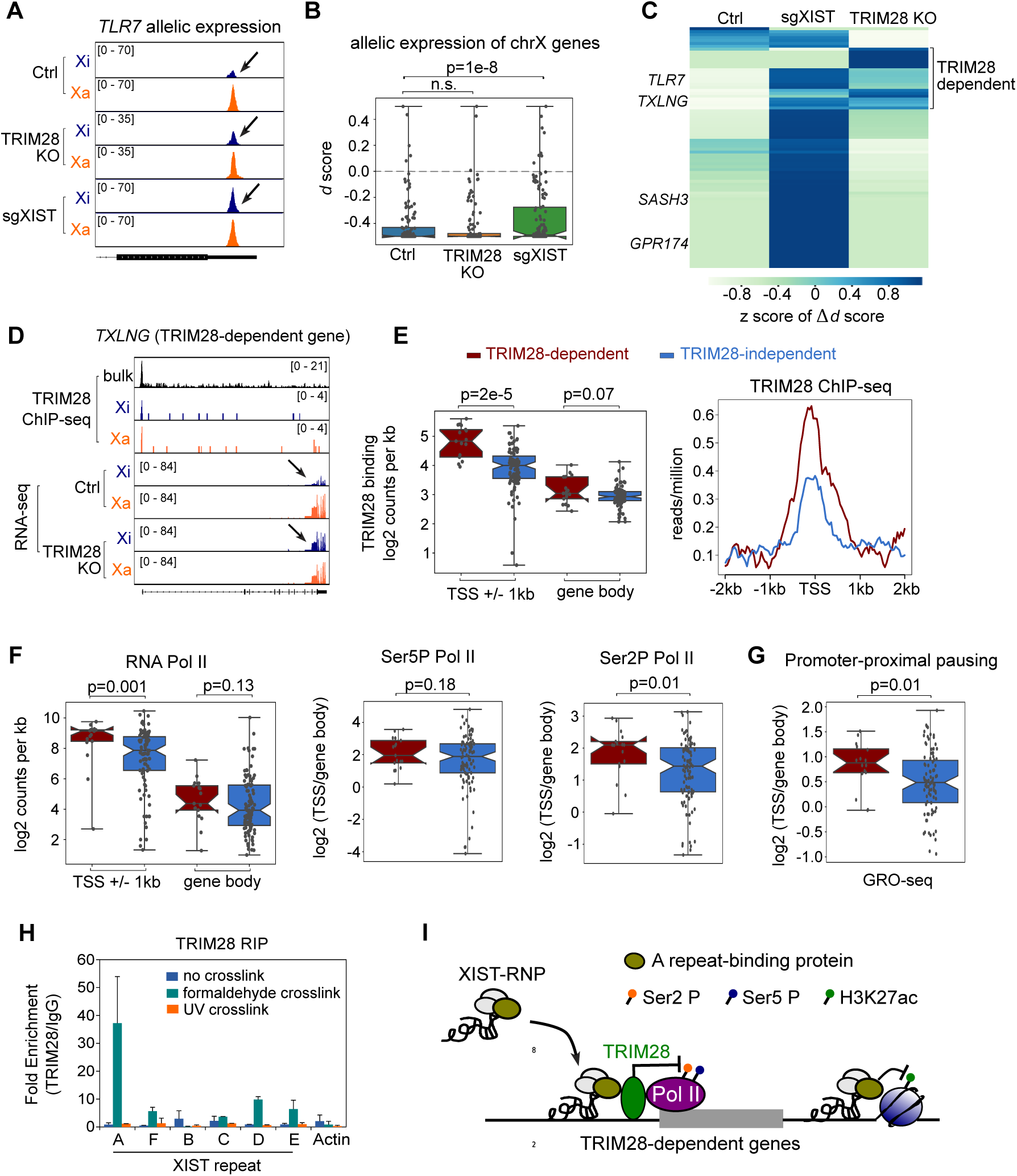
XIST cofactor TRIM28 participates in XCI maintenance of a subset of genes. A. Allelic RNA-seq track of *TLR7* gene at Xi (blue) and Xa (yellow) between control, TRIM28 KO and sgXIST group. B. Box plot showing the d score of chrX genes between control, TRIM28 KO and sgXIST group. P value was calculated by paired t-test. C. Heatmap showing the hierarchical clustering of genes with differential allelic expression between groups. Value depicts the z score of difference of the d score. D. Genome tracks showing TRIM28 ChIP-seq and RNA-seq in control and TRIM28 KO cells for representative TRIM28-dependent gene *TXLNG*. E. Box plot showing TRIM28 binding density (log2 counts per kb) at TSS+/-1kb or gene body regions between TRIM28-dependent and -independent genes (left). Average diagram showing the reads per million for TRIM28 ChIP-seq at TRIM28-dependent (red) and -independent genes (blue) at TSS +/-2kb (right). F. Box plot (left) showing RNA Pol II binding density (log2 counts per kb) at TRIM28- dependent (red) and -independent genes (blue). Box plots showing promoter pausing ratio (log2(read density^TSS^/read density^gene body^)) of Ser5 phosphorylated RNA Pol II (middle), and Ser2 phosphorylated RNA Pol II (right) at TRIM28-dependent (red) and -independent genes (blue). G. Box plot showing GRO-seq analysis of Pol II promoter pausing ratio (log2(read density^TSS^/read density^gene body^)) at TRIM28-dependent (red) and -independent genes (blue). H. Bar plot showing the TRIM28 RIP-qPCR on A-F repeats regions of XIST and Actin with no crosslink (blue), formaldehyde crosslink (green), and UV crosslink (orange). I. Schematic view of a proposed model showing TRIM28-mediated XCI maintenance. P values in E-G were calculated using non-parametric Mann-Whitney test.

Allelic analysis of TRIM28 ChIP-seq data revealed that TRIM28 binds to both Xa and Xi (**Figure S6D**). To confirm if the increased TRIM28 occupancy at TRIM28-dependent genes is not due to more binding at the Xa, we compared the allelic bias of TRIM28 ChIP-seq signal between TRIM28-dependent and -independent genes, and found a significantly increased binding at Xi rather than Xa of TRIM28-dependent genes compared to -independent genes (*p*=0.0008, Wilcoxon rank-sum test) (**Figure S6E**). Thus, unlike SPEN or Polycomb proteins that are enriched on the Xi (Dossin et al., 2020; Schoeftner et al., 2006), biallelic TRIM28 occupancy of a subset of X-linked genes can be modulated by XIST to maintain their X-inactivation.

TRIM28 is a transcriptional co-repressor protein that can evoke several silencing mechanisms. These include recruitment of SETDB1, a histone methyltransferase responsible for the deposition of the heterochromatin mark H3K9me3, interaction with HDAC-NuRD complex for histone deacetylation, or inhibiton of pTEFb to impact RNA polymerase pause release (Ma et al., 2019; Schultz et al., 2001, 2002). Therefore, we examined diagnostic epigenomic features of each of these mechanisms in turn. We observed variable H3K9me3 deposition between TRIM28-dependent (*TLR7*, *CXORF21*) and -independent (*BTK*, *ATP11C*) immune associated X-linked genes (**Figure S6F**). Systemically, we observed a higher H3K27ac deposition at promoters of TRIM28-dependent genes and no significant difference of repressive marks including H3K9me3 and H3K27me3 (**Figure S6G**), suggesting an alternative regulatory mechanism of TRIM28 for XCI maintenance that does not involve erasing H3K27ac or writing H3K9me3. TRIM28 is an E3 ligase that relies on RING domain and PHD domain. The PHD domain functions as intramolecular SUMO E3 ligase that is important for adjacent Bromodomain (BR) sumoylation, which facilitates recruitment of SETDB1 and NURD complex (reviewed in Czerwińska et al., 2017; Iyengar and Farnham, 2011). The RING domain functions as intermolecular SUMO E3 ligase that is responsible for CDK9 sumoylation (Ma et al., 2019). TRIM28 RING domain-mediated sumoylation inhibits CDK9 kinase activity, which is essential to release promoter-paused RNA polymerase II (PoI II) via Ser2 phosphorylation and thus block efficient transcription elongation (Ma et al., 2019). Given the potential regulatory role of TRIM28 in transcription elongation, we compared the binding profile of RNA Pol II, Pol II Ser5 phosphorylation (Ser5P, transcription initiation mark), and Pol II Ser2 phosphorylation (Ser2P, transcription elongation mark) between TRIM28- dependent and -independent genes. We found substantially increased Pol II occupancy at promoters of TRIM28-dependent genes, which is consistent with higher deposition of active mark H3K27ac (**Figure 6F, Figure S6G**). Interestingly, Ser2P enrichment at the promoter relative to the gene body is significantly higher in TRIM28-dependent genes compared to TRIM28-independent genes, whereas their Ser5P occupancy levels are quite similar (**Figure 6F**). This result indicates that TRIM28-dependent genes may possess higher promoter-proximal pausing of Ser2P Pol II and thus less efficient transcription elongation.

We further compared the Pol II promoter-proximal pausing ratio between TRIM28- dependent and -independent genes using GRO-seq data (Core et al., 2014). Indeed, there is a dramatic enrichment of nascent RNA at the promoter of TRIM28-dependent gene *TLR7*, while the nascent RNA is evenly distributed across the gene body for a TRIM28-independent gene, *SASH3* (**Figure S6H**). The GRO-seq analysis showed a significantly higher promoter-proximal pausing for TRIM28-dependent genes, suggesting that TRIM28 occupancy may inhibit transcription elongation of these genes (**Figure 6G**). Our data suggests that the XCI maintenance relies on TRIM28-dependent CDK9 sumoylation and subsequent Pol II pausing. To confirm whether the RING domain mediated CDK9 sumolyation instead of PHD domain mediated SETDB1 recruitment participates in XCI maintenance of TRIM28-dependent genes, we re-introduced TRIM28 full length protein or TRIM28 mutants with RING domain deletion or PHD-BR (PB) domain deletion into TRIM28 KO cells. We also created SETDB1 CRISPRi cells to test if TRIM28-mediated silencing is dependent on SETDB1. Indel analysis showed >80% knockout of endogenous TRIM28 in all groups (**Figure S6I**). The SETDB1 CRISPRi in B cells led to 82% knockdown of SETDB1 (**Figure S6J**). Allelic analysis by pyrosequencing on three TRIM28-dependent immune genes (*TLR7, CXORF21,* and *CXORF38*) showed that full length TRIM28 protein rescued the TRIM28 KO for gene silencing, as expected. Deletion of TRIM28 RING domain led to significant reactivation of all three TRIM28-dependent genes, suggesting RING domain-mediated CDK9 sumoylation and Pol II pausing are important for XCI maintenance of TRIM28-dependent genes. In contrast, deletion of TRIM28 PB domain or SETDB1 depletion had little effect on the allelic bias, indicating PB domain mediated SETDB1 recruitment and SETDB1-mediated H3K9me3 do not participate in silencing of X-linked TRIM28-dependent genes in B cells (**Figure S6K**). This finding is consistent with our result showing no significant difference of H3K9me3 deposition between TRIM28-dependent and -independent genes (**Figure S6G**).

Next, we performed TRIM28 RIP-qRT-PCR with formaldehyde crosslinking or UV crosslinking to assess the domain of XIST RNA that interacts with TRIM28 (**Figure 6H**). TRIM28 immunoprecipitation with formaldehyde crosslinking, but not UV crosslinking, specifically retrieved endogenous XIST A-repeat but not other portions of the lncRNA, suggesting that TRIM28 indirectly interacts with XIST possibly through interaction of XIST A-repeat binding proteins (**Figure 6H-I**).

Finally, to test if TRIM28 is required in XCI maintenance in a B cell-specific manner, we analyzed the published shRNA screen data in differentiated ES cells with inducible XIST expression and a GFP reporter allele (Moindrot et al., 2015). We found that TRIM28 shRNA is not enriched in GFP^hi^ cells compared to the input (**Figure S6L**), suggesting that TRIM28 is not required for X-inactivation in differentiated ES cells. SPEN shRNA is significantly enriched in GFP^hi^ cells, which is consistent with the well-established role of SPEN in XCI initiation during ES cell differentiation (Chu et al., 2015; McHugh et al., 2015). We also performed TRIM28 CRISPRi in K562 myeloid cells and achieved ∼70% knockdown of TRIM28 (**Figure S6M**). qRT-PCR analysis of showed that the loss of TRIM28 lead to a significant upregulation of four TRIM28-dependent X-linked genes only in B cells but not in K562 myeloid cells (**Figure S6K**). ChIP-seq analysis showed TRIM28 occupied these gene loci only in B cells but not in K562 myeloid cells (**Figure S6N**). Thus, both functional and biochemical evidence indicate cell-type specific action of TRIM28 on Xi genes in B cells.

Taken together, these results suggest that TRIM28 occupies the promoters of a subset of genes in B cells on both X chromosomes. Expression of XIST on the Xi and RNA-guided spread of the associated XIST binding proteins may allow these core factors to collaborate with TRIM28 on the Xi, and the TRIM28 RING domain-mediated sumoylation may prevent efficient transcription elongation of target genes to maintain their X-inactivation (**Figure 6I**).

### Escape of XIST-dependent genes in CD11c+ atypical B cells

It has been long speculated that defects in XCI maintenance may be associated with female-susceptible diseases including SS, SLE, and RA. Increased expression of X-linked genes along with the aberrant XIST localization was recently observed in immune cells from patients with lupus, suggesting abnormal XCI maintenance may predispose women to autoimmune diseases (Syrett et al., 2019). To delineate the biological function of XIST-mediated XCI maintenance, we performed functional annotation analysis on genes that are overexpressed after XIST depletion in human B cell line GM12878. The result showed significant enrichment of genes related to IFN-gamma cytokine production, and female-biased autoimmune diseases SLE and RA (**Figure 7A**), suggesting that XIST-mediated XCI maintenance may be essential to prevent overexpression of X-linked genes which may contribute to autoimmune diseases. To further test this idea, we used gene set enrichment analysis (GSEA) to assess the concordance of genes that are upregulated after the loss of XIST (including both autosome and X-linked genes) and genes activated in peripheral blood mononuclear cells (PBMCs) of patients with SLE over healthy donors. Importantly, we found that genes normally silenced by XIST tend to be over-expressed in patients with SLE compared to healthy donors such as genes *TLR7*, *CXCL10*, *IFI27*, *IFIT3*, *IFI6* and *OAS1* (**Figure S7A**). Recent studies showed that specific B cells such as IgD-CD27-CD11c+ double negative atypical B cells that are induced by unregulated TLR7 contribute to pathogenic responses in SLE and the TLR7 hyper-responsiveness may result in decreased B cell tolerance to self-antigen (Jenks et al., 2018). Our GSEA result showed that XIST-silenced genes tend to be over-expressed in these atypical B cells of lupus patients compared to healthy donors (**Figure S7A**). To further explore the impact of XIST-mediated XCI maintenance in SLE, we leveraged our discovery of “XIST-dependent” genes (X-linked genes whose maintenance are dependent on XIST) and “XIST-independent” genes (control X-linked genes whose maintenance are not). The GSEA analysis showed that XIST-dependent genes are significantly overexpressed in PBMC and IgD-CD27-CD11c+ atypical B cells of female SLE patients compared to healthy control, while the XIST-independent Xi genes were not (**Figure 7B, Figure S7B**). These results highlight the potential role of XIST dysregulation and subsequent escape of XIST-dependent genes in specific B cell subsets (CD11c+ atypical B cells) in certain diseases.

**Figure 7.**
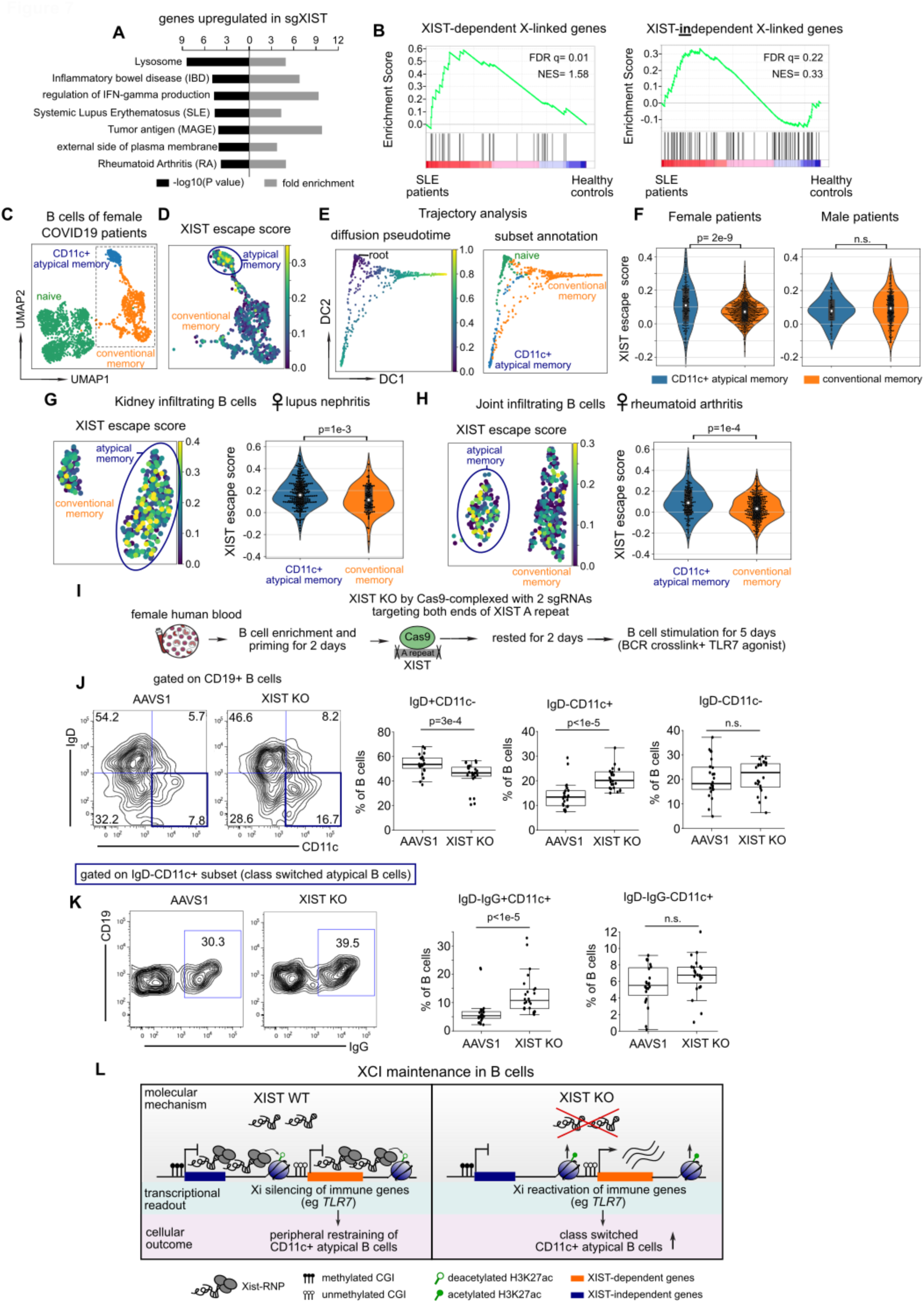
Escape of XIST-dependent genes in CD11c+ atypical B cells, which are increased after the loss of XIST. A. Bar plot showing significantly enriched biological pathway and diseases from genes that are upregulated after XIST perturbation in B cell line. B. GSEA analysis showing enrichment of XIST-dependent X-linked genes and XIST- independent X-linked genes in PBMC of female SLE patients compared to healthy donors. Normalized Enrichment Score (NES) and FDR q value are shown on the plot. C. Uniform manifold approximation and projection (UMAP) plot showing B cell subsets from female COVID-19 patients. D. UMAP plot showing XIST escape score at single cell level between CD11c+ atypical memory B cells and conventional memory B cells. E. Diffusion map showing the diffusion pseudotime trajectory via assigning naive cells as root (left). B cell subset projected on diffusion map (right). F. Violin plot showing the distribution of XIST escape score between CD11c+ atypical memory B cells and conventional memory B cells in female (lef) and male (right) COVID-19 patients. P value was calculated using unpaired t-test. G. UMAP showing XIST escape score of CD11c+ atypical memory cells and conventional memory B cells in kidney of female patients with lupus nephritis (left). Violin plot showing the the distribution of XIST escape score between CD11c+ atypical memory B cells and conventional memory B cells (right). P value was calculated using unpaired t-test. H. Same plot as in G of single cells from joint tissue of female patients with rheumatoid arthritis. I. Schematic view of deletion of XIST repeat A by Cas9-RNP electroporation in human primary B cells during B cell activation in response to BCR crosslinking and TLR7 ligand stimulation. J. Flow cytometry analysis of viable CD19+ human primary B cells between AAVS1 control and XIST KO group using IgD and CD11c markers (left). Box plot showing the frequency of IgD+CD11c-, IgD-CD11c+, IgD-CD11c-B cells among total B cells in two to three replicates from 8 female donors. P value was calculated using paired t-test. K. Flow cytometry analysis of IgG+ B cells gated on IgD-CD11c+ B cell subset. Box plot showing the frequency of IgD-IgG+CD11c+ and IgD-IgG-CD11c+ B cells among total B cells. P value was calculated using paired t-test. L. A proposed model of XCI maintenance in human B cells.

We developed a single cell gene expression score, termed “XIST escape score”, to evaluate XIST dysregulation in single cell transcriptome data across diverse human disease states and distinct B cell phenotypes. The XIST escape score is the normalized mean expression value of the XIST-dependent X-linked gene set minus that of the XIST-independent gene set. Thus, the score reflects the specific and coordinate induction of XIST-dependent genes while controlling for overall X-linked gene expression. Applying the XIST escape score to eleven human immune cell scRNA-seq data sets, we discovered specific disease states and B cell subsets where XIST dysregulation occur in vivo (**Table S7**). Interestingly, we found that escape of XIST-dependent genes occurs during COVID-19 infection, specifically in CD11c+ atypical memory B cells (**Figure 7C-D**). Single cell analysis distinguished CD11c+ atypical memory B cells from naive B and conventional memory cells with expression of a combination of specific markers (IgD-IgG+CD27midCD11c+T-bet+Zeb2+) (**Figure S7C-D**). Cell fate trajectory analysis revealed a bifurcation of the differentiation trajectory from naive B cells to conventional memory B cells vs. CD11c+ atypical B cells (**Figure 7E**). This observation is consistent with previous reports that CD11c+ atypical memory B cells exhibit a distinct differentiation fate with hyper-responsiveness to innate stimuli such as TLR7 and extra-follicular activation and expansion, compared to conventional memory B cells that are activated in germinal centers (Jenks et al., 2018; Tipton et al., 2015). Naive B cells with top 10 percentile of XIST escape score project strongly into atypical B cell fate (**Figure S7E**), suggesting that XIST dysregulation may bias this differentiation choice. XIST dysregulation, as reflected by the gain in XIST escape score, is evident in CD11c+ atypical B cells from female patients but not male patients (**Figure 7F**), consistent with this being a female-specific phenomenon related to X chromosome dosage. XIST dyregulation in atypical B cells during COVID-19 infection was replicated in two additional independent patient cohorts (**Table S7**).

We discovered a similar XIST dysregulation in kidney infiltrating B cells of lupus nephritis and joint tissue infiltrating B cells of rheumatoid arthritis in female patients. In both bases, CD11c+ atypical B cells demonstrate significantly increased XIST escape score compared to conventional memory B cells (**Figure 7G-H**). Importantly, this phenomenon is not observed in B cells from healthy control donors, nor female B cells in influenza viral infection, SFTS viral infection, or tumor-infiltrating B cells in breast cancer patients (**Table S7**). These results suggest that escape of XIST-dependent genes occurs in vivo in several diseases that associated with RNP antigens, and occurs specifically in CD11c+ atypical B cells that infiltrate the pathologic tissues.

Guided by our single cell trajectory analysis (**Figure S7E**), we hypothesized that XIST dysregulation may promote the differentiation of naive B cells to CD11c+ atypical memory cells. To test this, we deleted XIST in primary human B cells isolated from healthy female donors and monitored the B cell differentiation in response to BCR crosslinking and TLR7 stimulation that mimic viral single-strand RNA or excess autoreactive RNP stimulation. First, we utilized electroporation of Cas9-RNP complex (Cas9 protein assembled with 2 sgRNAs targeting both ends of the XIST A repeat) to knockout XIST in primary female B cells (**Figure 7I, Figure S7F**). To rule out the effect of DNA damage induced by Cas9 cutting, we designed control sgRNAs targeting a safe-harbor locus (AAVS1) to delete a fragment with similar length as XIST A-repeat. We found that the XIST A-repeat deletion led to ∼60% decreased expression of XIST RNA compared to the AAVS1 control (**Figure S7G**). After gating on viable CD19+ B cells (**Figure S7H**), we measured the frequency of class-switched CD11c+ atypical B cells (IgD-CD11c+) and found a significant increase of IgD-CD11c+ atypical B cells in XIST KO group compared to AAVS1 group (**Figure 7J**). It has been shown that TLR7 upregulation can drive the accumulation of CD11c+ atypical B cells (Jenks et al., 2018; Rubtsov et al., 2011). We speculated that the loss of XIST facilitates the overexpression of XIST-dependent immune gene TLR7 in primary B cells, which may lead to the increased frequency of CD11c+ atypical B cells. As expected, XIST deletion in primary human B cells showed ∼2.8 fold increase of TLR7 expression compared to AAVS1 deletion control (**Figure S7I**). TLR7 escape from XCI has been reported to associate with increased IgG class-switching in human B cells (Souyris et al., 2018). Indeed, we observed a significant increase of IgG class-switched CD11c+ atypical B cells after the loss of XIST (**Figure 7K**). Our single cell analysis of CD11c+ atypical B cells showed unique expression pattern including a mixture of IgD-CD27- and IgD- CD27+ populations and higher expression of transcription factors T-bet and Zeb2 compared to conventional memory cells (**Figure S7D**). To confirm if the loss of XIST lead to such features of CD11c+ atypical B cells upon B cell differentiation, we focused on the combination of IgD, CD27, and CD11c markers. Indeed, we found that both IgD-CD27-CD11c+ and IgD-CD27+CD11c+ B cell subsets are significantly increased in XIST KO group compared to AAVS1 control (**Figure S7J-K**). The qRT-PCR result also showed that the loss of XIST lead to increased expression of T-bet and Zeb2, key transcription factors for CD11c+ atypical B cells (**Figure S7L**).

In summary, based on the discovery of XIST-dependent genes in adult B cell line, we developed a computational strategy to track XIST dysregulation in any B cell gene expression data with single cell resolution, documenting the in vivo occurrence of escape of XIST-dependent genes in CD11c+ atypical B cells. We further experimentally validated that the loss of XIST can facilitate the naive cell differentiation into CD11c+ atypical B cells, a subset provide a first causal link between a known female-enriched B cell population (Rubtsov et al., 2011) with XIST dysregulation.

## DISCUSSION

### Continual requirement for XIST in XCI maintenance of somatic cells

Our study identified a central role of XIST in maintaining X-inactivation of genes in adult human B cells. Classic studies using somatic cell hybrids that deleted the X inactivation center first suggested that XIST is dispensable for XCI maintenance for a handful of X-linked genes examined (Brown and Willard, 1994). More recent studies using cultured cells with normal karyotypes and conditional knockout mouse models suggest a more complex picture, with a range of Xi gene reactivation and phenotypic consequences upon *Xist* removal (Adrianse et al., 2018; Bhatnagar et al., 2014; Csankovszki et al., 1999; Yang et al., 2016, 2020; Yildirim et al., 2013). These differences may be due to cell type-specific regulatory mechanism of XCI maintenance, and variable dependency of different X-linked genes on XIST for XCI maintenance. In contrast to mouse models that remove *Xist* in the context of development and are subject to compensation and selection (Yang et al., 2016, 2020), here we use CRISPR-based gene editing in adult human cells to achieve acute inactivation of XIST post development. We observed clear and robust escape of select genes from transcriptional repression on the Xi upon XIST inactivation. We note that B cells with *XIST* inactivation grows noticeably more slowly, and would be taken over by wild type cells with intact *XIST* in bulk culture over time, highlighting the need to perturb and observe in near synchrony. Because XCI is an epigenetic mechanism of gene memory over cell divisions, the human model with its substantially longer life span (max >100 years vs. 3 years for a mouse) and therefore far greater number of cell divisions provides a different scale of challenge for epigenetic memory. Targeting human XIST has recently been proposed as a potential therapeutic strategy to reactivate wild type genes on the Xi for female patients suffering from genetic disorders (Carrette et al., 2018). Thus, understanding XIST function in human somatic cells promises both biomedical gains and is technically feasible with modern epigenome editing technologies.

X-linked genes lacking promoter DNA methylation are particularly sensitive to XIST loss in B cells. These genes, escaped from XIST-dependent silencing when XIST is absent, can be re-silenced when XIST expression is restored. In addition, they are enriched in immune function, require continuous XIST RNP-mediated histone deacetylation in somatic B cells (**Figure 7L**). After blocking DNA methylation and H3K27me3 via small molecule inhibitors, genes classified as XIST-independent become reliant on XIST for XCI maintenance. These data demonstrate that the absence of DNA methylation or other chromatin-based memory makes XIST RNA-based transcriptional repression necessary. Thus, our data suggest a redundant model for XCI maintenance: epigenetic memory such as DNA methylation and XIST RNA-based transcriptional silencing can compensate each other to maintain a robust and durable X-inactivation. This is in line with a previous study revealing a synergism of XIST RNA and DNA methylation in XCI maintenance (Csankovszki et al., 2001). This model further suggests that the classification of “XIST-dependent” vs “XIST-independent” may be fluid and cell-type specific; the target gene behavior may be predicted based on the level of DNA methylation and other chromatin marks at the locus in the relevant cell type. Our expanded discovery of XIST requirement in other somatic cell types including fibroblasts and neuronal cells highlight the ongoing requirement for XIST can be generalized although with important differences in scale.

### Cell type-specific XIST RNPs to prevent escape from X-inactivation

The XIST interactome is comprised of shared and distinct protein partners in ES cells, myeloid cells, and B cells. We leveraged the XIST ChIRP-MS and allelic CRISPRi screen to functionally annotate the XIST interactome in XCI maintenance in adult B cells. The major silencing mechanism of XIST appears to be histone deacetylation, as evidenced by SPEN, the A-repeat binding adaptor protein, being the top scoring CRIPSRi hit, and validated by allelic RNA-seq in SPEN RRM2-4 deletion B cells. The N-terminus of SPEN contains 4 RRM domains that directly interact with the A-repeat (Lu et al., 2016) while the C-terminal SPOC domain recruits and activates histone deacetylase activity of HDAC3 (Dossin et al., 2020; Żylicz et al., 2019). CRISPRi of XIST tethering proteins HNRNPU or CIZ1 individually has little effect on XCI maintenance, suggesting a redundant role for these two proteins or alternative co-factors for XIST tethering in B cells such as YY1 (Wang et al., 2016). E repeat binding proteins PTBP1 and MATR3 are essential for XCI maintenance in B cells, suggesting a key role of E repeat in maintaining X-inactivation. Interestingly, a recent study showed that deletion of E repeat has no effect on XIST coating during onset of XCI but leads to dispersed localization of XIST and the loss of Xi silencing at a later stage (Pandya-Jones et al., 2020). Such continuous silencing is dependent on the formation of protein-condensate via interaction with PTBP1 and MATR3, connecting phase separation with XCI maintenance (Pandya-Jones et al., 2020).

Our data revealed TRIM28 as a B-cell specific XIST cofactor that is critical to XCI maintenance of a subset of X-linked genes. Since allelic binding of TRIM28 is not enriched at the Xi and there is no direct interaction between TRIM28 and XIST based on UV RIP, we reason that XIST does not directly recruit TRIM28 to the Xi. It is possible that when XIST recruit proteins to spread across the Xi, the XIST RNP comes into contact with TRIM28 that pre-bound at certain loci, leading to indirect interaction between TRIM28 and XIST RNP on chromatin that we detected by CHIRP-MS and RIP experiments with formaldehyde crosslinking (**Figure 6I**). TRIM28 interacts specifically with the A-repeat of XIST, suggesting potential functional collaboration with other silencing factors, such as SPEN and RBM15 that bind to the A-repeat. Our data suggest two models by which TRIM28 acts to promote XCI maintenance: (1) Functionally complementation to block different steps of target gene transcription. TRIM28 acts at promoters while SPEN and other XIST-associated factors acts on preferentially on enhancers (Dossin et al., 2020; Żylicz et al., 2019). (2) Activity modification: XIST-RNP may stimulate TRIM28 silencing function through modifying its activity such as SUMO E3 ligase activity. 35% of TRIM28-dependent genes are not dependent on XIST for silencing, favoring the model of functional cooperation rather than the model of activity modification. We showed that TRIM28 promoter occupancy is significantly associated with RNA Pol II promoter-proximal pause to cooperate with XIST-mediated H3K27 deacetylation at enhancers, unveiling a novel regulatory mechanism for XCI maintenance (**Figure 6I**). Finally, Carter et al. recently proposed a model of X inactivation by viral mimicry, based on the discovery that XIST A-F repeats evolved from insertion of endogenous retroviruses (ERV) and that SPEN, the key XCI factor, has an evolutionarily ancient role in binding ERV RNAs to elicit transcriptional silencing across autosomes (Carter et al., 2020). It is well established that TRIM28 is a major restriction factor to silence ERVs via chromatin modification (Fasching et al., 2015), providing support for the idea that the machinery for ERV silencing may be coopted for dosage compensation. TRIM28 also has major role in the regulation of B cell genes and B cell development (de Sio et al., 2012), which is suitable for its cooption in B cell XCI maintenance.

The discovery of unique XIST RNPs in somatic cells suggest that different somatic cells can have overlapping but still different XIST co-factors mediated silencing to prevent XCI escape. Like a transcription factor that collaborates with additional transcription coactivators and corepressors to regulate different target genes in different cell types, the same lncRNA can also recruit, assemble, and guide different protein complexes to enact distinct consequences in different cell types. Thus, future work on identification of cell type-specific XIST binding proteins may provide insights into sex-biases in diseases such as autoimmunity, inflammatory, neurodegenerative and cardiovascular disorders (Möller-Leimkühler, 2007; Skuse, 2005). Our computational analysis of XIST dysregulation tracking using existing single-cell transcriptome data and experimental validation using CRSIPR-Cas9 mediated deletion of XIST in human primary cells can be a powerful tool to dissect the mechanism and function of XCI maintenance in primary cell types that are critical to sex dimorphism in human health and diseases.

### XIST dysregulation in pathogenic atypical B cells in human disease

We developed XIST escape score to track XIST dysregulation at single cell resolution in an unbiased manner in vivo and across diverse disease states. We discovered that XIST dysregulation, reflected by escape of XIST-dependent genes, occurs in CD11c+ atypical B cells in patients with female-biased autoimmune diseases including SLE and RA, and ssRNA viral infection such as COVID-19. Sex is a major determinant of clinical outcomes in COVID-19 and drives different immune responses in men vs. women (Takahashi et al., 2020; Williamson et al., 2020). We further showed that the XIST dysregulation introduced by XIST KO in female primary B cells, facilitates the formation of CD11c+ atypical B cells. Consistently, this unique B cell subset exhibits a strong female bias only in aged mice where the hormone difference is minimized, further suggesting that XIST-mediated XCI maintenance may contribute to the sex dimorphism of this subset (Rubtsov et al., 2011; Rubtsova et al., 2015). Our data showed that the deletion of *XIST* results in higher expression of X-linked immune gene *TLR7*, and in the context of BCR ligation and TLR7 stimulation, increased the frequency of CD11c+ atypical B cells, a subset that is highly dependent on TLR7 signal for their activation and formation. Interestingly, our functional annotation of XIST-silenced genes showed enrichment of the production of IFN-gamma, a key cytokine that has been shown to induce the expression of T-bet, a master transcription factor that promotes the differentiation of CD11c+ atypical B cells upon TLR7 stimulation (Zumaquero et al., 2019). Indeed, we observed increased expression of T-bet after the loss of XIST. Therefore, our results suggest that higher expression of TLR7 and T-bet after the loss of XIST may increase the responsiveness to TLR7 ligand stimulation, thus providing a selective advantage under TLR7 agonist to select out the CD11c+ atypical B cells. In addition to TLR7, other X-linked immune genes may also participate into the sex dimorphism. For example, we found the interferon inducible X-linked gene CXORF21 as an XIST-dependent gene (**Figure 1F**), and CXORF21 is encompassed in a risk haplotype significantly associated with SLE, a female-biased autoimmune disease (Odhams et al., 2019). CXORF21, also known as TASL, has recently been discovered as a key adaptor protein for endolysosomal TLR7 to activate downstream transcription (Heinz et al., 2020). Therefore, XIST is essential to balance the gene dosage of X-linked immune genes that can greatly impact the the formation of CD11c+ atypical B cells, a rare population in healthy donors but aberrantly expanded in certain infectious diseases and female-biased autoimmune diseases (Cancro, 2020). A recent study observed an absence of XIST localization to the Xi in CD11c+ atypical B cells but not conventional memory B cells (Pyfrom et al., 2020), consistent with our functional studies. Future work on how XCI maintenance is impaired in this subset will provide a better understanding of sex dimorphism in CD11c+ aypitcal B cells.

## STAR METHODS

### KEY RESOURCES TABLE

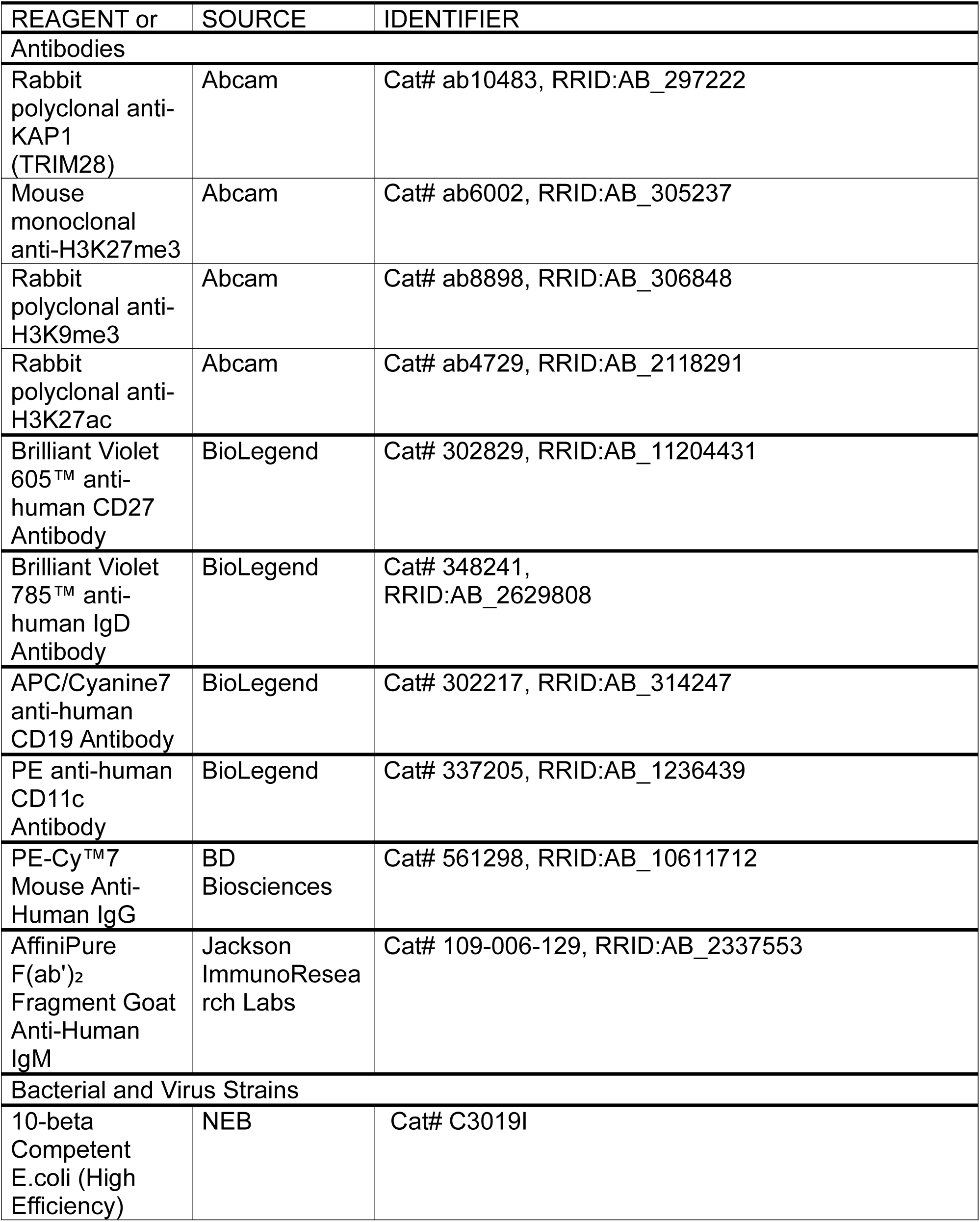
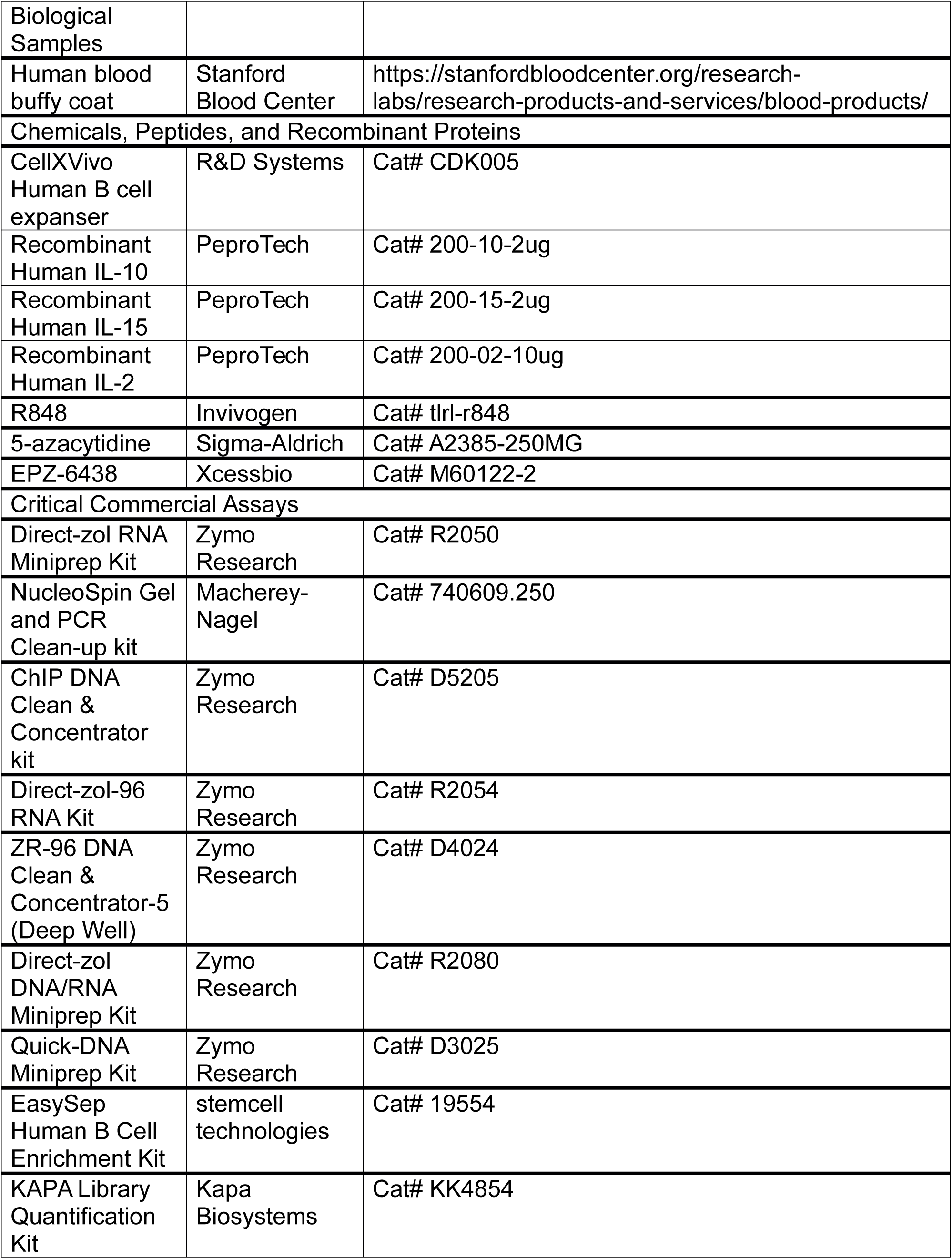
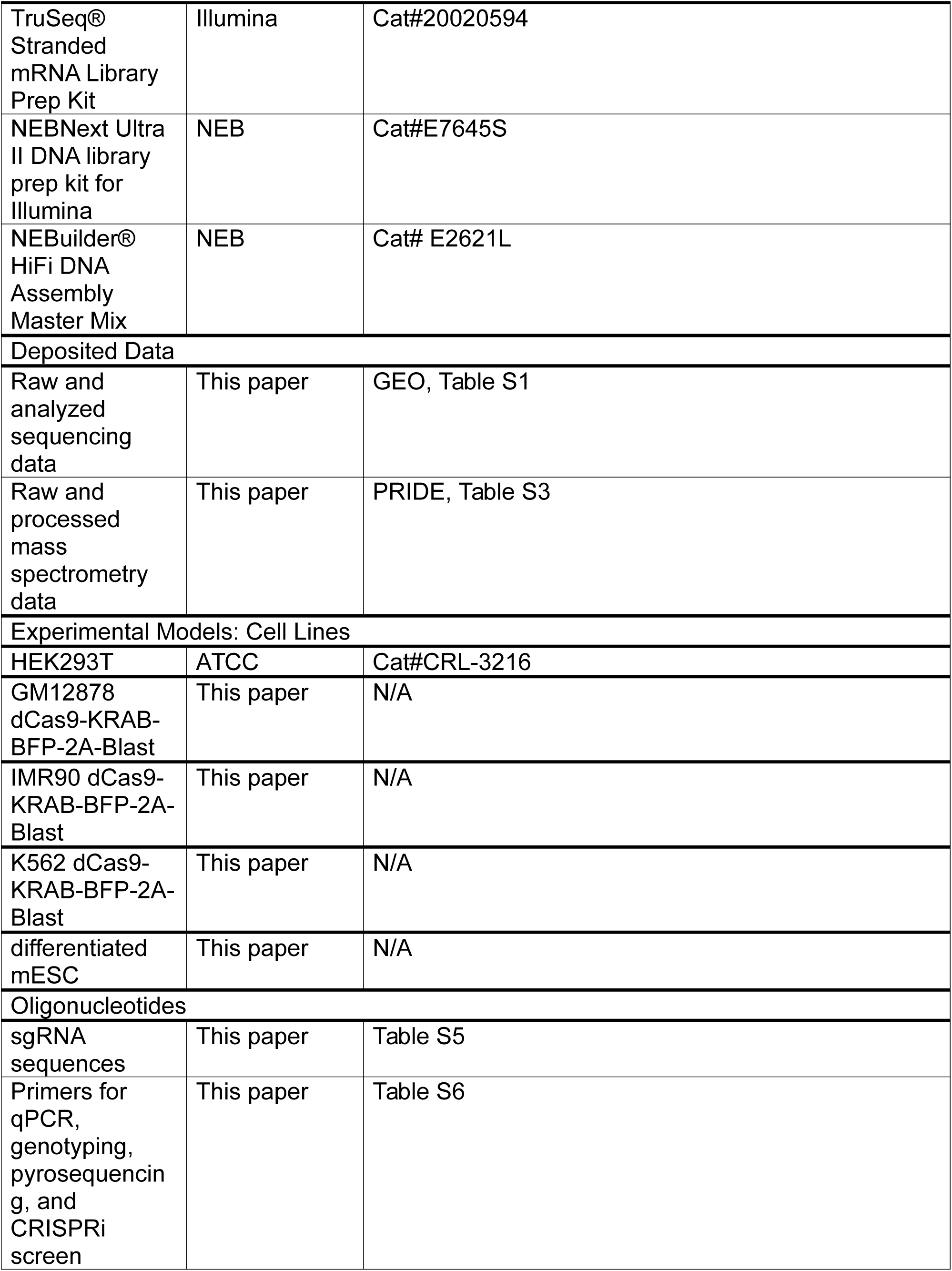
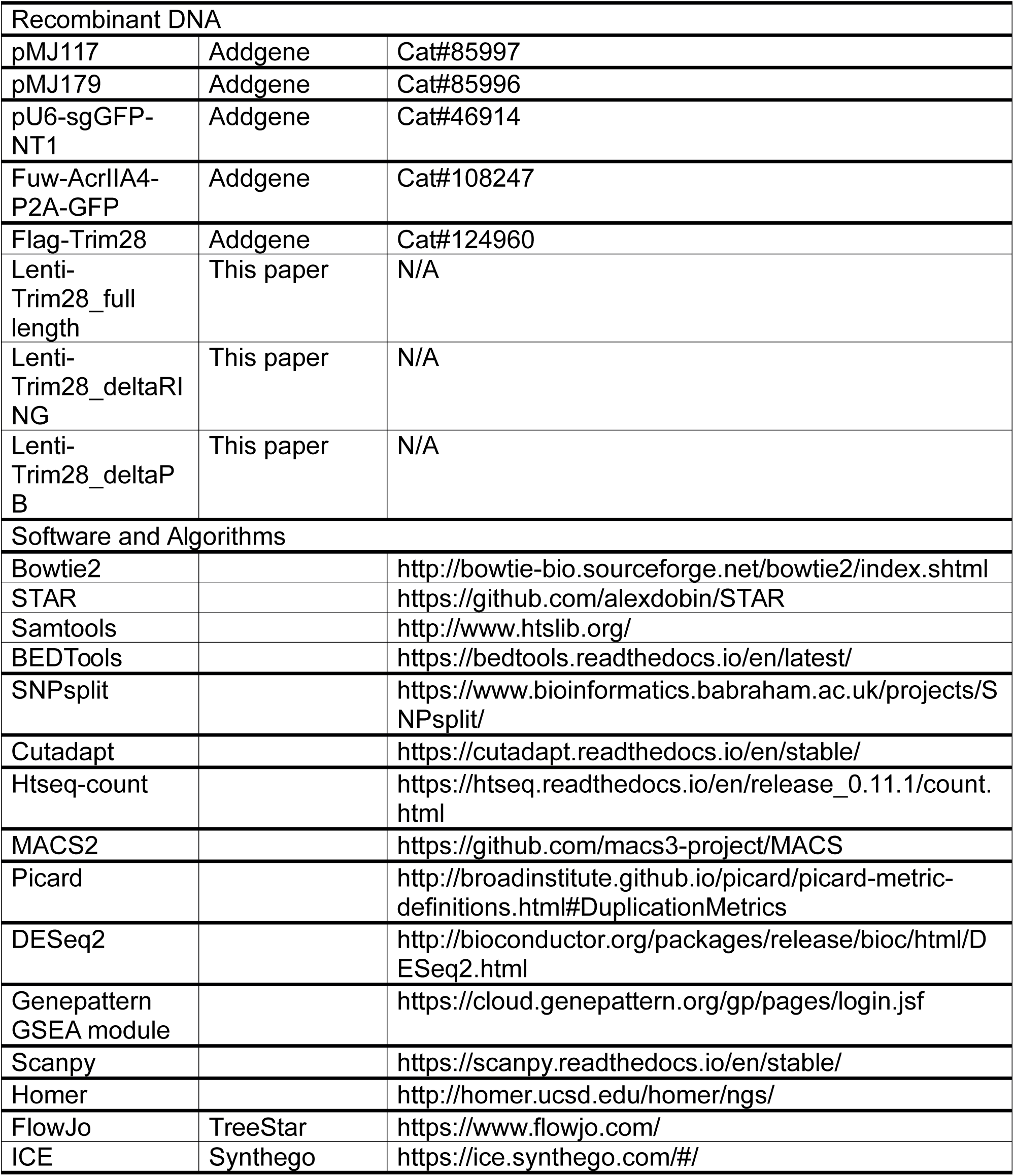

### CONTAC FOR REAGENT AND RESOURCE SHARING

Further information and requests for resources and reagents should be directed to and will be fulfilled by the Lead Contact, Howard Y. Chang.

### EXPERIMENTAL MODEL AND SUBJECT DETAILS

#### Generation of stable cell lines and cell culture

GM and K562 cell lines were cultured in RPMI 1640 (Thermo Fisher Scientific) with 10% FBS (Thermo Fisher Scientific) and 1% Penicillin-Streptomycin (Gibco). HEK 293T cells were cultured in DMEM (Thermo Fisher Scientific) with 10% FBS and 1% Penicillin-Streptomycin. IMR90 cell line were cultured in EMEM (ATCC) with 10% FBS and 1% Penicillin-Streptomycin. Female mESC were cultured on gelatin-coated plates in N2B27 medium for 7 days and then differentiated into NPC using N2B27 medium supplemented with EGF and FGF (both 10ng/mL, Peprotech). To inhibit DNA methylation and H3K27me3, B cell line was treated with 0.3 uM DNMT inhibitor 5-azacytidine and 2 uM EZH2 inhibitor EPZ-6438 for 7 days.

For generation of XIST CRISPRi GM B cell line, we performed lentiviral transduction of sgXIST in dCas9-KRAB expressing GM B cell line which has been previously made in the lab (Rubin et al., 2019). We designed two sgRNAs targeting the XIST promoter and cloned them into one lentiviral vector as previously described (Rubin et al., 2019). To generate the CRISPRi virus, we cultured HEK 293T cells at 4 million per 10cm dish and transfected with 4.5 ug pMP.G, 1.5 ug psPAX2 and 6 ug sgRNA vector using OptiMEM and Lipofectamin 3000 (Cat# L3000015, Thermo Fisher Scientific) at the following day. Two days later, the supernatant was collected and filtered with 0.44 um filter. The virus was concentrated 1:10 using Lenti-X Concentrator (Clontech). dCas9-KRAB+ GM cells were seeded at 300K cells per well of a 6-well plate and 40 uL concentrated virus was added to the media. Two days later, we select the XIST sgRNA expressing cells by adding 2 ug/mL puromycin for seven days and cells expressing mCherry were sorted by BD Aria II FACS sorter. SETDB1 CRISPRi GM B cell line is generated similarly as XIST CRISPRi GM B cell line.

For generation of anti-CRISPR expressing XIST CRISPRi or control CRISPRi B cell line, we performed lentiviral transduction of anti-CRISPR protein in non-targeting control or XIST (sgXIST) CRISPRi GM B cell line which is generated above. To generate the anti-CRISPR virus, we cultured HEK 293T cells at 4 million per 10cm dish and transfected with 4.5 ug pMP.G, 1.5 ug psPAX2 and 6 ug Fuw-AcrIIA4-P2A-GFP vector (Addgene #108247) using OptiMEM and Lipofectamin 3000 at the following day. Two days later, the supernatant was collected and filtered with 0.44 um filter. The virus was concentrated 1:10 using Lenti-X Concentrator. SgNT or sgXIST CRISPRi cells were seeded at 300K cells per well of a 6-well plate and 40 uL concentrated virus was added to the media. A week later, we select the anti-CRISPR and CRISPRi expressing cells by sorting cells expressing both GFP and mCherry using BD Aria II FACS sorter. The anti-CRISPR efficiency is measured by qRT-PCR of XIST.

For generation of XIST CRISPRi IMR90 cell line, we first established dCas9-KRAB expressing IMR90 cell line by lenti-viral transduction of dCas9-BFP-KRAB (Addgene #46911) and sorted BFP+ cells by BD Aria II FACS sorter. Next, we performed transduction of lentivirus expressing sgXIST or sgNT (non-targeting control) in dCas9-KRAB expressing IMR90 cell line and selected by adding 2 ug/mL puromycin for 7 days and sorted cells expressing both BFP and mCherry. The CRISPRi efficiency is measured by qRT-PCR of XIST.

For generation of TRIM28 CRISPRi K562 cell line, we first established dCas9-KRAB expressing K562 cell line by lenti-viral transduction of dCas9-BFP-KRAB and sorted BFP+ cells by BD Aria II FACS sorter. Next, we performed transduction of lentivirus expressing sgTRIM28 or sgNT in dCas9-KRAB expressing K562 cell line and sorted cells expressing both BFP and mCherry. The CRISPRi efficiency is measured by qRT-PCR of TRIM28.

For generation of XIST A repeat KO GM B cell line, we used CRISPR Cas9 RNP editing to cut XIST A repeat. SgRNAs were produced by combining tracr and crRNA according to the Synthego Biosciences protocol. Two sgRNAs (30 uM) targeting both ends of XIST repeat A were complexed with 3 ug Cas9 protein (IDT) for 10 min and then electroporated to GM cells (1700V, 20ms, 1 pulse) using Neon Transfection System (Thermo Fisher Scientific). Transfected cells were sorted into 96-well plate for single cell per well and then single cell clone with repeat A deletion was screened by genomic DNA PCR and RT-qPCR.

GM single cell clones with SPEN RRM2-4 deletion were generated similarly as XIST A repeat KO and screened by genomic DNA PCR and RT-qPCR. GM single cell clones with TRIM28 deletion were generated similarly and screened by western blot. All sgRNA sequences were listed in **Table S5**.

### METHOD DETAILS

#### CHIRP-MS and proteomics analysis

25 T152 flasks of GM12878 or K562 cells were used for ChIRP-MS (0.5-1 billion cells) as previously described (Chu et al., 2015). Cells were cross-linked in 3% formaldehyde for 30 min followed by 0.125 M glycine quenching for 5 min. Cross-linked cells were lysed in fresh NLB buffer (50 mM Tris pH 7.0, 10 mM EDTA, and 1% SDS) and sonicated in 1 mL Covaris tube for 20 min with the following parameters (Fill level:10; Duty Cycle:15; PIP:140; Cycles/burst:200). The supernatant was pre-cleared with streptavidin beads (Invitrogen) and then treated with or without 30 ug/mL Rnase A (Cat # EN0531, Thermo Fisher Scientific) at 37°C for 45 min. Then the supernatant was hybridized with 8.8 uL XIST probes in Hybridization Buffer (50 mM Tris pH 7.0, 1 mM EDTA, 1% SDS, 750 mM NaCl, and 15% Formamide) at 37°C rotating overnight and added with 880uL of streptavidin beads and rotate for 45 min at 37°C. Beads were washed 5 times in ChIRP Wash Buffer (2X SSC and 0.5% SDS) for 5 min at 37°C. RNA extraction can be performed using a small aliquot of post-CHIRP beads to assess the XIST RNA retrieval efficiency. XIST bound proteins were eluted from beads in Biotin Elution Buffer (12.5 mM biotin (Invitrogen), 7.5 mM HEPES, 75 mM NaCl, 1.5 mM EDTA, 0.15% SDS, 0.075% sarkosyl, 15% Formamide, and 0.02% Na-Deoxycholate) at RT for 20 min then shake at 65°C for 10min, finally precipitated by TCA at 4°C overnight. The proteins were pelleted at 16000 rcf at 4°C for 30 min and the protein pellets were washed once with cold acetone and air-dry for 1 min, then the proteins were solubilized in 1X laemmli sample buffer and boiled at 95C for 30 min. Final proteins were size-selected on 4-12% NuPAGE gels for mass spectrometry. Gel slices were excised and diced to 1 mm cubes prior to proteolytic digestion. Samples were reduced in 5 mM DTT in a 50 mM ammonium bicarbonate buffer at 55°C for 30 mins. After removing the residual solvent, proteins were alkylated with 10 mM acrylamide in the same buffer, rinsed with 50% acetonitrile, and then digested using Trypsin-Lys C (Promega) overnight at 37°C to obtain peptides. Samples were centrifuged to condense particulates so that the solvent including peptides could be collected. A further peptide extraction was performed by the addition of 60% acetonitrile, 39.9% water, 0.1% formic acid and incubation for 10-15 min before collection. Samples were dried by speed vac prior to resuspension in 2% aqueous acetonitrile with 0.1% formic acid for mass spectrometry. Mass spectrometry experiments were performed on an Orbitrap Fusion Tribrid mass spectrometer (Thermo Scientific, San Jose, CA) with liquid chromatography using an Acquity M-Class UPLC (Waters Corporation, Milford, MA). For a typical LC/MS experiment, a flow rate of 450 nL/min was used, where mobile phase A was 0.2% formic acid in water and mobile phase B was 0.2% formic acid in acetonitrile. Analytical columns were pulled and packed in-house using fused silica with an I.D. of 100 microns packed with Dr. Maisch 1.8 micron C18 stationary phase to a length of ∼25 cm. Peptides were directly injected onto the analytical column using a gradient (2-45% B, followed by a high-B wash) of 80min. The mass spectrometer was operated in a data-dependent fashion using CID fragmentation for MS/MS spectra generation collected in the ion trap. The collected mass spectra were analyzed using Byonic (Protein Metrics) for peptide identification and protein inference. The search was performed against the Uniprot *homo sapiens* database, including isoforms. Cysteine modified with propionamide was set as a fixed modification in the search, with other typical modifications, *e.g.* oxidation of methionine, included as variable modifications. .Data were held to a 12 ppm mass tolerance for precursors and 0.4 Da for MS/MS fragments, allowing up to two missed cleavage sites. Data were validated using the standard reverse-decoy technique at a 1% false discovery rate. The peptide counts in experiment group and Rnase treatment group for each replicate in GM and K562 cells are listed in **Table S3**. The XIST binding proteins were first identified in each replicate with peptide counts >25 and fold enrichment (experiment group/Rnase treatment group) >1.5. Then the proteins that are overlapped between replicates were identified and further filtered using mean fold enrichment from two replicates>2.5. The highly-confident XIST binding proteins for CRISPRi screen were identified using peptide counts>25 and mean fold enrichment >10.

#### RNA immunoprecipitation (UV-RIP and formaldehyde-RIP)

The RNA immunoprecipitation protocol was performed as previously described ((G Hendrickson et al., 2016) with a few modifications. For UV-RIP, 5 million cells per replicate were crosslinked by UV exposure at 0.3 J/cm2 (254nM UV-C). For formaldehyde-RIP (fRIP), 5 million cells per replicate were fixed in 0.1% formaldehyde for 10 min at RT and quenched by 0.125 M glycine for 10 min at RT. UV-crosslinked or formaldehyde-fixed cells were washed by cold PBS and lysed in lysis buffer (50 mM Tris, 150 mM Kcl, 0,1% SDS, 1% Triton-X, 5 mM EDTA, 0.5% sodium deoxycholate, 0.5 mM DTT, 1XPIC, 100 U/mL RNAseOUT inhibitor) and sonicated using a Covaris Ultrasonicator tube (140W, 10% duty cycle, 200cycles/burst). After 15 min max speed spin at 4°C, the supernatant was pre-cleared with Protein A/G beads for 30 min at 4°C and the beads were disposed. An aliquot of the supernatant was saved as the total input control. The leftover supernatant was incubated with 2 ug mouse IgG (Thermo Fisher) or 2 ug KAP1 (ab10483, Abcam) and rotated overnight at 4°C. 20ul Protein A/G beads were added and rotated for 2 h at 4°C and washed four times with native lysis buffer (25 mM Tris, 150 mM KCl, 5 mM EDTA, 0.5% NP-40, 1X PIC, 100 U/mL RNAseOUT inhibitor, 0.5 mM DTT). The IP beads and input samples were reverse-crosslinked in RCL buffer (2% N-lauroyl Sarcosine, 10 mM EDTA, 5 mM DTT in PBS without Mg and Ca) with Proteinase K and RNAseOUT inhibitor for 1h at 42°C and then another 1h at 50°C, shaking at 1000 rpm. The RNA was purified using Quick-RNA Miniprep Kit (Zymo Research). The RNA quality was checked by BioAnalyzer and the qRT-PCR was performed using Stratagene Brilliant II SYBR Green QRT-PCR Master Mix (Agilent). The primers for RIP qPCR were listed in **Table S6**.

#### RNA extraction, RT-qPCR, and pyrosequencing

RNA was extracted in TRIzol using Quick-RNA Miniprep Kit following the manufacturer’s protocol with on-column Dnase digestion (Zymo Research). RT-qPCR was performed using Stratagene Brilliant II SYBR Green QRT-PCR Master Mix (Agilent). To quantify the allelic bias, cDNA was generated by reverse transcription of extracted RNA using SuperScriptIII and the amplified for 5 cycles using targeted qPCR primers. Then the PCR product was purified by Zymo DNA Clean & Concentrator-5 and further amplified using biotinylated primers and sequenced on using Q24 Pyromark (Qiagen). All primers for qRT-PCR and pyrosequencing are listed in **Table S6**.

#### RNA-seq

RNA was extracted using Quick-RNA Miniprep Kit with on-column Dnase digestion (Zymo Research). At least 100ng RNA was used to prepare the RNA-seq library using TruSeq® Stranded mRNA Library Prep Kit (Cat# 20020594, Illumina) for each sample following the manufacturer’s instruction. The library was sequenced on an Illumina Hiseq 4000 to generate 2X75 paired-end reads or 2X150 paired-end reads.

#### Rescue experiment with TRIM28 mutants

Since TRIM28 KO B cells are lethal, we developed a strategy to first express TRIM28 mutants and then knock out the endogenous TRIM28. Lentiviral vectors that carry full length TRIM28 and RING domain or PB (PHD-Bromodomain) domain deletion mutants conjugated to eGFP fused with a 2A peptide were cloned using Gibson Assembly from the PCR product of the plasmid containing TRIM28 cDNA (Addgene#). The N-terminal 180 bp sequence of all TRIM28 variants was altered to distinguish with endogenous TRIM28 sequence with silent mutation (producing same amino acids with endogenous TRIM28). Lentivirus expressing TRIM28 mutants were made similarly as above. B cells were transduced with TRIM28 mutants expressing lentivirus for 5 days and GFP^hi^ cells were sorted for endogenous TRIM28 knockout. Two sgRNAs (30 uM) targeting the sequence inside of the N-terminal 180 bp of endogenous TRIM28 were complexed with 3 ug Cas9 protein (IDT) for 10 min and then electroporated to GFP^hi^ B cells (1700V, 20ms, 1 pulse) using Neon Transfection System. The knockout efficiency was measured by indel from sanger sequencing of genomic DNA PCR product of the first 180bp. The percentage of indels was calculated by ICE analysis (Synthego).

#### CUT&RUN

CUT&RUN for H3K27me3 and H3K9me3 were performed as previously described in (Skene and Henikoff, 2017). Briefly, 0.5 million cells per replicate were bound to 10 uL concanavalin A-coated beads (Cat# BP531, Bangs Laboratories) in Binding Buffer (20 mM HEPES, 10 mM KCl, 1 mM CaCl2 and 1 mM MnCl2). The beads were washed and resuspended in Dig-Wash Buffer (20 mM HEPES, 150 mM NaCl, 0.5 mM Spermidine, 0.05% Digitonin, and 1X PIC). The primary antibody of H3K27me3 (Ab6002, Abcam, 1ug) or H3K9me3 (Ab8898, Abcam, 1ug) was added to the bead slurry and rotated at 4°C overnight. The beads were washed by Dig-Wash Buffer and the secondary antibody (1:50 for rabbit anti-mouse IgG) was added and rotated at 4°C for 1 h, and washed by Dig-Wash buffer. The pA-Mnase fusion protein (a gift from Henikoff lab) was added at 700ng/mL and rotated at 4°C for 1 h. Next, the beads were washed by Dig-Wash buffer twice followed by Low-Salt Rinse Buffer (50 mM HEPES and 0.02% Digitonin). The targeted digestion was performed by incubating at 0°C in Incubation Buffer (3.5 mM HEPES, 10 mM CaCl2, and 0.02% Digitonin) for 30 min and stopped by 1X STOP buffer (20 mM EGTA, 0.02% Digitonin, 25 ug/mL RnaseA, 50 ug/mL Glycogen, and 170mM NaCl). The chromatin was released by incubating at 37°C for 30 min. The DNA extraction was performed by NucleoSpin Gel and PCR Clean-up kit (Cat# 740609.250, Machery-Nagel). The CUT&RUN library was prepared using NEBNext Ultra II DNA library prep kit for Illumina (NEB) following the manufacturer’s protocol, and finally sequenced on an Illumina Hiseq 4000 to generate 2X75 paired-end reads.

#### ChIP-seq

For H3K27ac ChIP-seq, 1 million cells per replicate were fixed in 1% formaldehyde for 10 min at RT and quenched by 0.125 M glycine for 10 min at RT. For Trim28 ChIP-seq, 1 million cells per replicate were first cross-linked with 2 mM DSG for 40 min at RT and then fixed in 1% formaldehyde for 10 min at RT and finally quenched by 0.125 M glycine for 10 min at RT. Fixed cells were first lysed in membrane lysis buffer (50 mM HEPES-KOH, 140 mM NaCl, 1 mM EDTA, 10% glycerol, 0.5% NP-40, 0.25% TritonX-100) for 10 min at 4°C. Then pelleted nuclei were lysed in nuclear lysis buffer (10mM Tris, 200mM NaCl, 1mM EDTA, 0.5mM EGTA) for 10min at RT. The chromatin was resuspended in Covaris sonication buffer (10 mM Tris, 1 mM EDTA, 0.1% SDS) and sonicated in a Covaris Ultrasonicator milliTube (140W, 10% duty cycle, 200 cycles/burst) to get 150-700bp length. The sonicated lysate was incubated with 4 ug H3K27ac (ab4729, Abcam) or 5 ug KAP1 (ab10483, Abcam) overnight at 4°C. Antibody-bound chromatin was incubated with Protein-G Dnyabeads (10004D, Invitrogen) for 4 h at 4°C and washed five times with RIPA wash buffer (5 mM HEPES-KOH, 500 mM LiCl, 1 mM EDTA, 1% NP40, 0.7% Sodium Deoxycholate) and eluted in elution buffer (10 mM Tris, 10 mM EDTA, 1% SDS). The eluted sample was incubated at 65°C overnight to reverse the crosslink and treated with RNAse A for 1h at 37°C and Proteinase K for 1h at 55°C. DNA was extracted using ChIP DNA Clean & Concentrator kit (Zymo Research). The ChIP-seq library was prepared using NEBNext Ultra II DNA library prep kit for Illumina (NEB) following the manufacturer’s protocol, and finally sequenced on an Illumina Hiseq 4000 to generate 2X 75 paired-end reads.

#### CRISPRi screen for allelic expression of TLR7

We designed two sgRNAs per XIST co-factor gene with each targeting a different region in 200 bp after the transcriptional start site using CRISPOR. One sgRNA each was cloned into pMJ117 (human U6 vector) or pMJ179 (mouse U6 vector) which were digested with BstXI and BlpI using NEBuilder HiFi DNA Assembly Master Mix (NEB). Then the corresponding U6 promoter and sgRNA sequences were PCR amplified and cleaned up by ZR-96 DNA Clean & Concentrator-5 (Deep Well). The PCR fragments were assembled into the pU6-sgGFP-NT1 lentiviral vector which was digested with XbaI and XhoI, using NEBuilder HiFi DNA Assembly Master Mix. Finally, the individual colonies with 2X sgRNA plasmid were screened by colony PCR and confirmed by Sanger sequencing. To generate CRISPRi lentivirus, we plated 800K HEK293T cells per well for 6-well pate and for the next day we transfected 0.75 ug pMP.G, 0.25 ug psPAX2, and 1 ug sgRNA vector in 100 uL Opti-MEM using Lipofectamin 3000. After two days, the supernatant was collected and the virus was concentrated 1:10 using Lenti-X Concentrator (Clontech). 200K dCas9-KRAB expressed GM cell line were plated per well in 24 well plates and 40 uL concentrated virus was added the following day. Two days later, we select the Xist-cofactor sgRNA expressing cells by adding 2 ug/mL puromycin for 2-3 weeks. Selection media was refreshed every three days. RNA was extracted using Direct-zol-96 RNA Kit (Zymo). TLR7 RT-PCR was performed using Stratagene Brilliant II SYBR Green QRT-PCR Master Mix (50°C for 30 min, 95°C for 10 min, then 35 cycles of 95°C for 30 seconds, 60°C for 1 min, and 72°C for 30 seconds, finally 72°C for 5 min). The second PCR with Ad1_TLR7_F and Ad2_TLR7_R primers was performed using NEBNext Master Mix (2.5uL 1^st^ PCR product with 500 nM primers in program: 98°C for 1 min, 20 cycles of 98°C for 30 seconds, 60°C for 30 seconds, and then 72°C for 40 seconds, finally 72°C for 5 min). The third PCR with indexed primers containing barcodes was performed using NEBNext Master Mix (2uL of 1:10 diluted 2^nd^ PCR product with 1uM indexed primers in program: 98°C for 1 min, 10 cycles of 98°C for 30 seconds, 72°C for 30 seconds, and then 72°C for 40 seconds, finally 72°C for 5 min). Finally, all the samples were pooled and purified using Zymo DNA clean & concentrator. The library was size-selected to 250-300 bp. The gel slices were cut and the DNA was recovered by Zymo Gel DNA recovery kit (Zymo Research). The concentration of the library was quantified by KAPA Library Quantification Kit for Illumina (Roche) and sequenced on an Illumina Miseq 2X75 cycle. Reads were mapped directly to TLR7 reference Xa and Xi allele and counted at each allele for d score analysis. The co-factors with d score^co-factor^ - d score^Ctrl^ >0.1 were labeled as essential XIST co-factors for TLR7 XCI maintenance.

#### Human primary B cell editing with Cas9 RNP

Buffy coats from female healthy donors were obtained from Stanford Blood Center with consent forms. Peripheral blood mononuclear cells (PBMC) were isolated using Lymphoprep (Cat# 07811, STEMCELL Technologies) density-gradient centrifugation and cryopreserved and stored in -80C. B cells were purified from thawed PBMCs by negative selection using EasySep Human B Cell Enrichment Kit (Cat#19844, STEMCELL Technologies) following the manufacturer’s protocol. Isolated B cells were cultured in IMDM medium supplemented with 10% FBS and 55 mM beta-mercaptoethanol at 1X10^6^ cell/mL and primed with CellXVivo Human B cell expander (1:250 dilution, R&D system), 50 ng/mL IL10 (Cat#200-10-2ug, PeproTech), 10 ng/mL IL15 (Cat#200-15-2ug, PeproTech), 50 ng/mL IL2 (Cat#200-02-10ug, PeproTech) for two days. For B cell electroporation, primed B cells were washed by PBS twice and resuspended in Neon Buffer T (Cat# MPK1025, Thermo Fisher Scientific). Two sgRNAs (30 uM) targeting both ends of XIST repeat A were complexed with 3 ug Cas9 protein (IDT) for 10 min at RT. Cas9 RNP was added to the resuspension to get the final cell density at 3 X10^7 cells/ml and the electroporation was performed (1700V, 20ms, 1pulse) in 10 uL Neon transfection system. Electroporated cells were immediately transferred to pre-warmed B cell expansion medium (RPMI 1640 +10% FBS +1XGlutaMAX (Thermo Fisher Scientific) + 50 ng/mL IL2 + CellXVivo Human B cell expander (1:250 dilution, R&D system)). After two days of culture, an aliquot of cells was saved to test editing efficiency. The left cells were activated in B cell expansion medium supplemented with 1 ug/mL TLR7 agonist R848 (Invivogen) and 5 ug/mL F’2 Fragment Goat Anti-Human IgM (Cat# C840J42, Jackson ImmunoResearch Laboratories) for 5 days and subjected to flow cytometry.

#### Flow cytometry

B cells were first incubated with Human TruStain FcXTM (Fc Block, BioLegend) in cell staining buffer (Biolegend, Cat# 420201) for 30 min on ice and then stained with viability dye (Thermo Fisher, Cat # R37610), CD27 (clone O323), IgD (clone IA6-2), IgG(clone G18-145), CD19 (clone HIB19), CD11c (clone Bu15) for 20 min on ice. The stained cells were washed in 1X FACS buffer for twice and they ready for flow cytometry. All flow cytometry antibodies are from Biolegend or BD Biosciences. All flow cytometry was performed using LSRII and FACS data were analyzed by FlowJo.

#### Computational analysis for RNA-seq

RNA-seq reads were mapped to the human genome (hg19) using STAR with default parameters (--outFilterMultimapNmax 1 --alignEndsType EndToEnd --outSAMattributes NH HI NM MD). Mitochodria reads were removed by samtools and duplicated reads were removed by picard. Quantification of aligned reads at the gene level was performed by HTseq count with default parameters (--stranded=reverse –additional-attr=gene_name). Raw counts were used to identify differentially expressed genes (DEG) using DESeq2 with size factor (total reads) normalization and DEGs were identified if Benjamini & Hochberg adjusted p-value <0.05. For Gene Ontology (GO) term analysis in Figure S1H, PANTHER GO was used. For gene functional annotation in Figure 7A, DAVID gene-annotation enrichment, KEGG pathway, and GAD disease category were used with cut-off (Benjamini & Hochberg adjusted p-value < 0.05 and fold enrichment > 3). In Figure 7, GSEA analysis was performed using Genepattern GSEA module with public available female SLE PBMC and B cell subset RNA-seq data (Jenks et al., 2018; Tokuyama et al., 2018). The gene sets used in GSEA analysis (genes upregulated in sgXIST, XIST-dependent X-linked genes, and XIST-independent X-linked genes) were listed in **Table S2**.

For allelic-specific RNA-seq analysis, SNP sites for donor NA12878 were extracted from the dbSNP database. SNP sites that are shared between paternal and maternal alleles but different from the human genome were kept as the common NA12878 SNP. SNP sites that are different between paternal and maternal allele were replaced by ‘N’ to build up N-masked reference genome. To maximize the allelic read depth, two biological replicates were pooled for later analysis. RNA-seq reads were aligned to the N-masked reference genome by STAR. After alignment and removal of mitochondria reads, remaining reads were split to maternal-specific (Xa) or paternal-specific (Xi) allele using SNPsplit. Duplicated reads were removed by picard and quantification of allele-specific reads at the gene level was performed via htseq-count similar to the bulk analysis. Genes with allelic reads >10 were retained to calculate the d score (reads^Xi^/(reads^Xa^+reads^Xi^)-0.5). For genes with d score <0, genes with d score^sgXIST^-d score^ctrl^ > 0.03 and binomial P value <0.05 are defined as XIST-dependent genes. The other genes are XIST-independent genes. X-linked immune gene set was curated from review (Libert et al., 2010). The hierarchical clustering was performed using seaborn clustermap with parameters (metric=euclidean, z_score=0). In Figure S1G, the variable escapee, constitutive escapee and the X-inactivated genes were defined from GTEx study (Tukiainen et al., 2017). In Figure S2C-D, the effect size of sex bias for variable escapees in different tissues and the category of variable escapee, constitutive escapee, and the X-inactivated genes were defined from GTEx study (Oliva et al., 2020). In Figure 6, genes with d score^TRIM28KO^-d score^ctrl^ > 0.03 and d score^ctrl^ <0 are defined as TRIM28-dependent genes. The other genes with d score^ctrl^ <0 are defined as TRIM28-independent genes. Bedgraph files were generated from bam files using bedtools genomcov and then convert to bigwig files for IGV visualization.

#### Computational analysis for CUT&RUN and ChIP-seq

Reads were mapped to the human genome (hg19) using bowtie2 with default parameters (--very-sensitive). Mitochondrial reads were removed by samtools and duplicated reads were removed by picard. For allelic-specific analysis, to maximize the allelic read depth, two biological replicates were pooled for later analysis. Reads were aligned to the N-masked reference genome by bowtie2. After alignment and removal of mitochondria reads and duplicated reads, remaining reads were split to Xa or Xi allele using SNPsplit. For ChIP-seq of H3K27ac, peaks were called using MACS2 with parameters (--broad -f BAMPE –broad-cutoff 0.01) over input control. The total consensus peak coordinate across samples was obtained using bedtools merge and intersect. The peaks were assigned to the nearest genes using HOMER annotatePeaks. The peaks within TSS +/- 1kb range were defined as promoters and other distal H3K27ac peaks were defined as enhancers. Allelic reads overlapping on peaks were counted using bedtools intersect and peaks with the allelic reads >10 were retained to calculate the d score (reads^Xi^/(reads^Xa^+reads^Xi^)-0.5). For H3K27me3 and H3K9me3 CUT&RUN and TRIM28 ChIP-seq, reads were aligned and filtered as above. TRIM28 ChIP-seq in K562 cell line is downloaded from public ENCODE dataset ENCFF165QYW. The windows were binned as TSS adjacent regions (TSS +/- 1kb) and gene bodies (TSS+1kb to TES) for each gene. Total and allelic reads overlapping these regions were counted using bedtools and regions with the allelic reads >10 were retained to calculate the d score. Average plot of TRIM28 bulk signal was generated by ngsplot. Bedgraph files were generated from bam files using bedtools genomcov and then convert to bigwig files for IGV visualization.

#### Feature comparison

ChIP-seq of H3K27ac (ENCFF180LKW), RNA Pol II (ENCFF368HBX), Ser5P RNA Pol II (ENCFF002UPS), and Ser2P RNA Pol II (ENCFF031RUV), MEDIP-seq (GSE56774), and GRO-seq (GSE60454) from public available ENCODE and GEO datasets were used for feature comparison. ATAC-seq of GM12878 cells (Buenrostro et al., 2013) and our CUT&RUN of H3K9me3 and H3K27me3 in control GM cells were used for feature comparison. Reads were aligned to hg19 or N-masked hg19 genome by bowtie2 with mitochondria reads and duplicated reads removal as above. The windows were binned as TSS adjacent regions (TSS +/- 1kb) and gene bodies (TSS+1kb to TES) for each gene. Total and allelic reads overlapping these regions were counted using bedtools and regions with the allelic reads >10 were retained to calculate the d score. GC content was calculated by bedtools nuc. CpG island, LINE, SINE, and LTR files were downloaded from UCSC Table browser (RepeatMasker) and their density at TSS adjacent regions or gene bodies were counted by bedtools. For linear regression analysis, counts of each feature across X-linked genes were scaled from 0 to 1 individually and analysis was performed by statsmodels linear regression with OLS model. For logistical regression analysis, train and test sets for features were split by sklearn train_test_split with test_size=0.25. Then analysis was performed using sklearn LogisticRegression. GRO-seq analysis was performed using HOMER. The promoter-proximal index was calculated as read density^TSS adjacent regions^/read density^gene bodies^ for each X-linked gene.

#### Computational analysis for single-cell RNA-seq

Public single-cell RNA-seq data were downloaded from GEO database (GSE149689, GSE154567, GSE155673, GSE142016, GSE159117, GSE149313, GSE145281, GSE135710) and ImmPort repository (SDY997, SDY998). The single cell data preprocess is done by Scanpy version 1.4.6 (Wolf et al., 2018). The sex of the sample is validated by the expression of male-specific gene DDX3Y, and female-specific gene XIST. The genes that are detected in less than 3 cells were removed using sc.pp.filter_genes(). The cells with fewer than 200 detected genes and higher than 10% mitochondrial genes were removed. The data was normalized to 10000 reads per cell for total-count to correct the library size using sc.pp.normalize_total() with target_sum=1e4. The data was then log transformed using sc.pp.log1p(). The top 2000 highly-variable genes were identified using sc.pp.highly_variable_genes() with min_mean=0.0125, max_mean=3, min_disp=0.5. The total counts per cell and the percentage of mitochondrial genes expressed were regressed out using sc.pp.regress_out(). Each gene is scaled to unit variance using sc.pp.scale() with max_value=10. Principle component analysis was performed using top 2000 highly variable genes using sc.tl.pca(). Then the top 40 principle components were used to compute the neighborhood graph for dimensional reduction using sc.pp.neighbors() with n_neighbors=10, n_pcs=40, random_state=1. The Leiden graph-clustering method was applied to find Leiden clusters using sc.tl.leiden() with different resolution based on datasets. The B cell clusters were subset based on B cell marker genes MS4A1 and CD79A while excluding other immune cell types based on cell-specific marker genes such as CD3E, CD14, PPBP, CD4, CD8A, FCER1A, and CST3 etc. The clusters for B cell subsets were manually annotated as naive, conventional memory, atypical memory B cells based on known cell type markers (IGHD, IGHG1, CD27, ITGAX). The UMAP plots on these marker genes for B cell subsets were generated by sc.tl.umap() and sc.pl.umap(). The compact violin plot for more marker genes of these B cell subsets was generated by sc.pl.stacked_violin(). For XIST escape score analysis, the score is calculated by mean expression of XIST-dependent X-linked genes subtract mean expression of XIST-independent X-linked genes (gene sets from Table S2). The calculation is performed using sc.tl.score_genes() with XIST-independent genes as reference gene pool and XIST-dependent genes as target gene list (ctrl_size=50, n_bins=25). The p value of gene score between subsets was calculated by unpaired t-test using scipy stats ttest_ind() function. For trajectory analysis, a weighted nearest neighbor graph was first constructed t0 build a transition matrix by sc.pp.neighbors() with n_neighbors=20, method=’gauss’. Then the naive B cell is assigned as a root cell. The diffusion map was computed using n_comps=15 by sc.tl.diffmap(). The Diffusion Pseudotime order was computed by sc.tl.dpt() with n_branchings=1, n_dcs=10 to identify branching point and differentiation endpoint. The diffusion pseudotime plot was generated by sc.pl.diffmap().

## ACKNOWLEDGEMENT

We thank CK. Chen, K.L. Huang, N. Pacalin, A. Carter, J. Xu, members of the Chang lab and E. Heard for discussion. We also thank R. Leib at Mass Spectometry core, PAN facility, FACS facility and SFGF sequencing core (S10OD018220) at Stanford University. Supported by Scleroderma Research Foundation, Pershing Square Foundation, NIH U54-CA260517, NIH RM1-HG007735 (to H.Y.C.), and a Stanford Dean’s Fellowship (to B.Y.). A.T.S. was supported by the NIH K08CA230188 and a Career Award for Medical Scientists from the Burroughs Wellcome Fund. H.Y.C. is an Investigator of the Howard Hughes Medical Institute.

## AUTHOR CONTRIBUTIONS

B.Y. and H.Y.C conceived the project. B.Y. performed most experiments and analyzed the data. Y.Q. and A.T.S. performed CHIRP-MS. R.L. performed RNA-seq library preparation. Q.S. performed TRIM28 mutants cloning. B.Y. and H.Y.C. wrote the manuscript with input from all authors. H.Y.C. supervised the work and acquired the funding.

## DISCLOSURE

H.Y.C. is a co-founder of Accent Therapeutics, Boundless Bio, and an advisor to 10x Genomics, Arsenal Biosciences, and Spring Discovery. A.T.S. is a co-founder of Immunai.

## SUPPLEMENTARY FIGURE LEGEND

**Figure S1.**
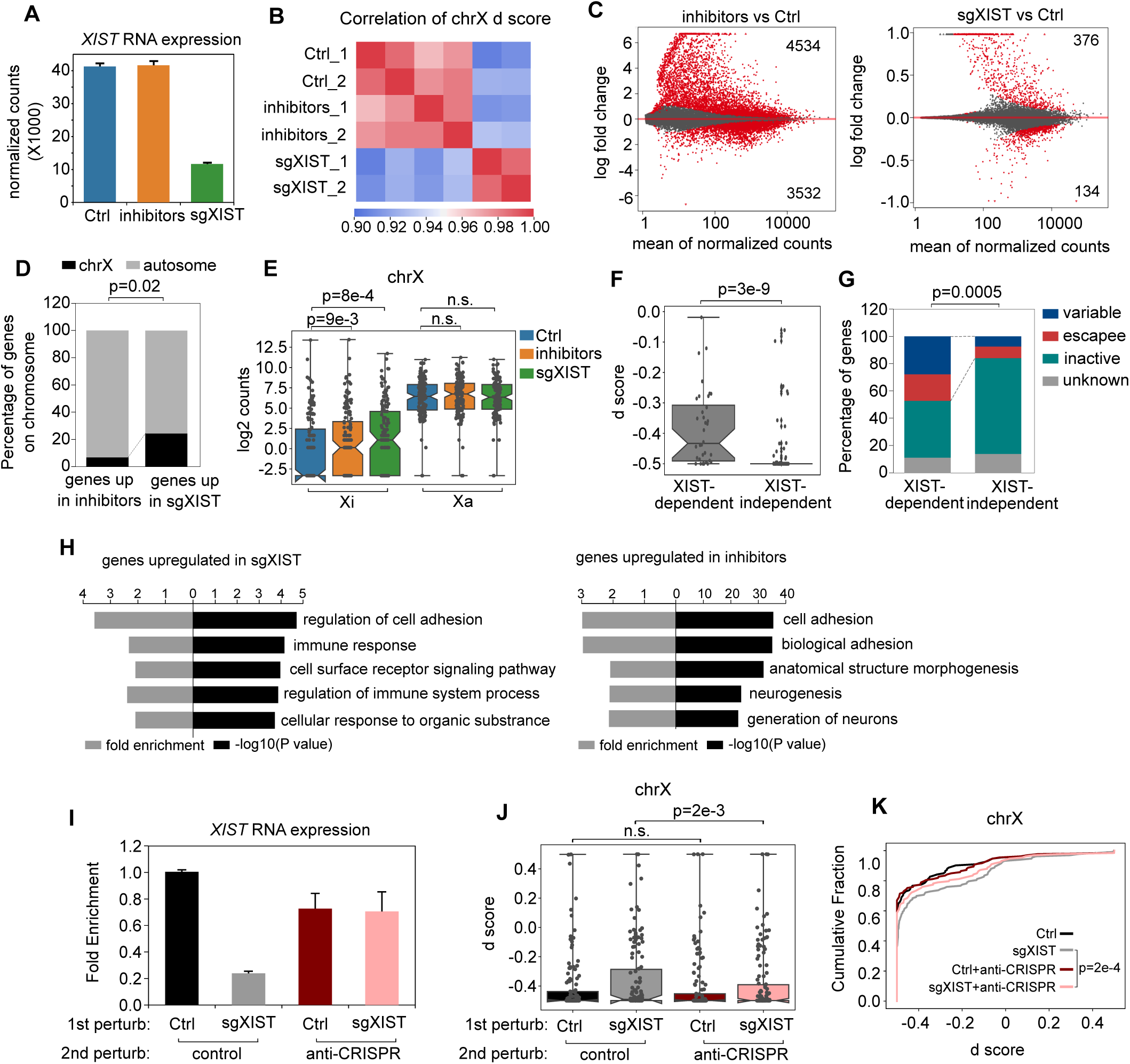
Allelic-specific RNA-seq in B cell line perturbed with XIST or treated with epigenetic inhibitors. Related to Figure 1. A. RNA-seq analysis showing the XIST RNA expression across control, sgXIST and inhibitors group. B. Heatmap showing Pearson correlation of gene expression allelic bias (d score) on chromosome X between replicates with the control group (Ctrl), XIST perturbation group (sgXIST) and inhibitors treatment group (inhibitors). C. MA plot showing differentially-expressed genes (highlighted in red) between control and inhibitors group (left), or between control and sgXIST group (right). Numbers showing significantly upregulated genes (top) or downregulated genes (down). D. Bar plot showing the percentage of genes on chrX and autosomes between genes that are upregulated in inhibitors group and genes that are upregulated in XIST group compared to control group. P value was calculated using Fisher exact test. E. Box plot showing the log2 counts of expression of genes across the chrX at Xi and Xa in ctrl, sgXIST and inhibitors group. P value was calculated using paired t-test. F. Box plot showing the d score of XIST-dependent and XIST-independent genes in control group using allelic RNA-seq data. P value was calculated using non parametric Mann-Whitney test. G. Bar plot showing the percentage of variable escapee, constitutive escapee and inactive genes between XIST-dependent and -independent genes. P value was calculated using Fisher exact test. H. Gene Ontology analysis showing top GO terms enriched in genes upregulated in sgXIST group (left) or genes upregulated in inhibitors group (right). I. qRT-PCR result of XIST RNA expression of B cells that have been transduced with sgXIST or non-targeting control and then overexpressed with or without anti-CRISPRI protein (ACRIIA4). J. Box plot showing the distribution of d score of allelic gene expression on the X chromosome in groups as in I. P value was calculated using paired t-test. K. Cumulative distribution of d score of allelic gene expression across the X chromosome in groups as in I. P value was calculated using Kolmogorov–Smirnov test.

**Figure S2.**
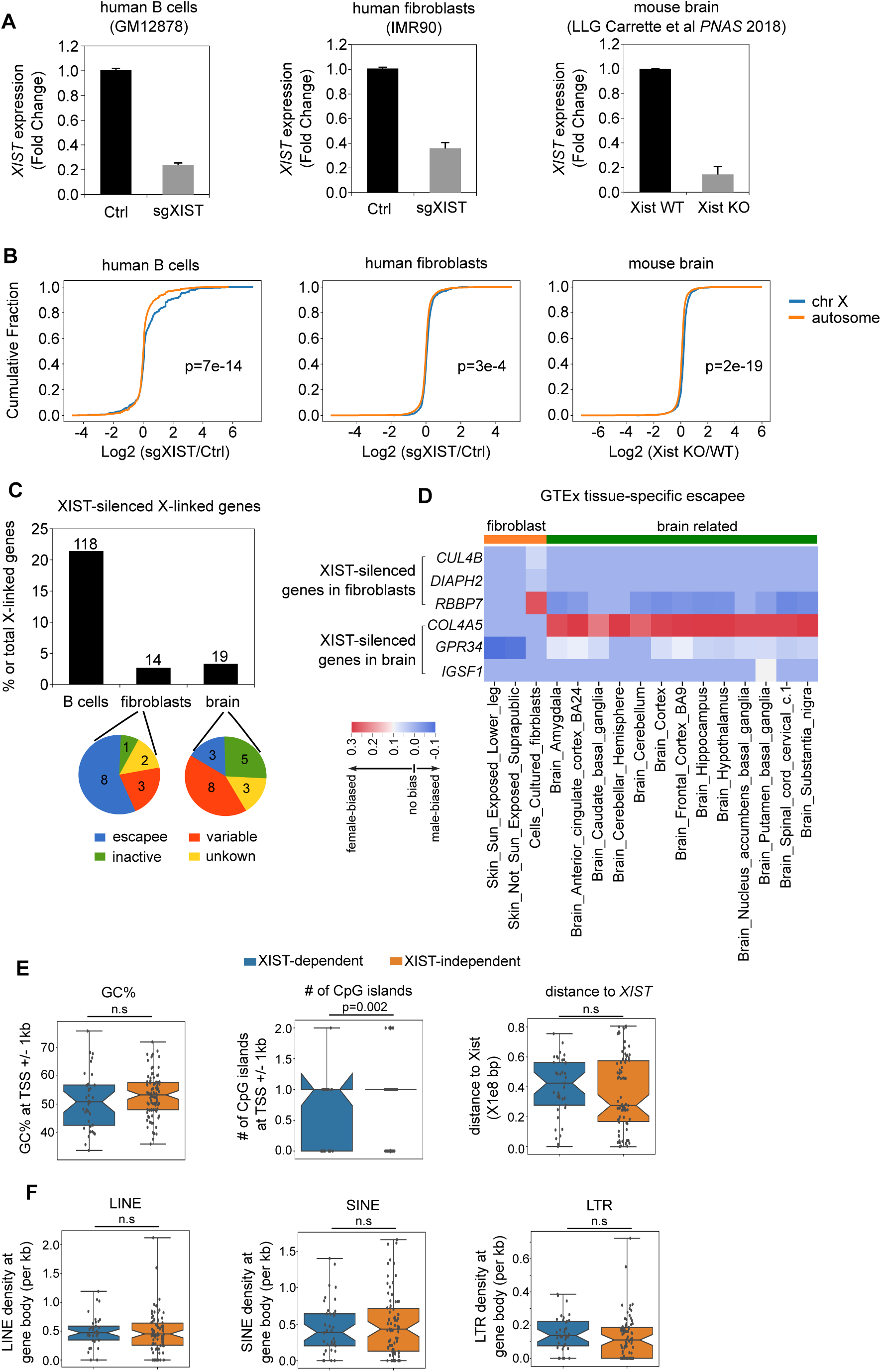
Comparison of genetic features between XIST-dependent and -independent genes. Related to Figure 2. A. Bar plot showing the relative gene expression of XIST from control and sgXIST group in female human B cell line (GM12878) and female human fibroblast cell line (IMR90) from qRT-PCR, or neuronal cells from Xist WT and brain-specific Xist KO mice from RNA-seq data. B. Cumulative distribution of log2(sgXIST/control) fold between X chromosome and autosomes in three cell types as in A. P value was calculated using Kolmogorov–Smirnov test. C. Bar plot showing the percentage of XIST-silenced X-linked genes in three cell types. Pie chart showing the number of escapee, variable escapee, X-inactivated genes, and unknown category defined from GTEx study among XIST-silenced X-linked genes in fibroblasts and neuronal cells. D. Heatmap showing effect size of sex bias in different tissues of variable escapees among XIST-silenced genes in fibroblasts and neuronal cells. E. Box plots showing the GC percentage, the number of CpG islands, and the distance to XIST between XIST-dependent (blue) and -independent (yellow) genes. P values were calculated via non-parametric Mann-Whitney test. F. Box plots showing the density of transposon element LINE, SINE and LTR between XIST-dependent (blue) and -independent (yellow) genes. P values were calculated by non-parametric Mann-Whitney test.

**Figure S3.**
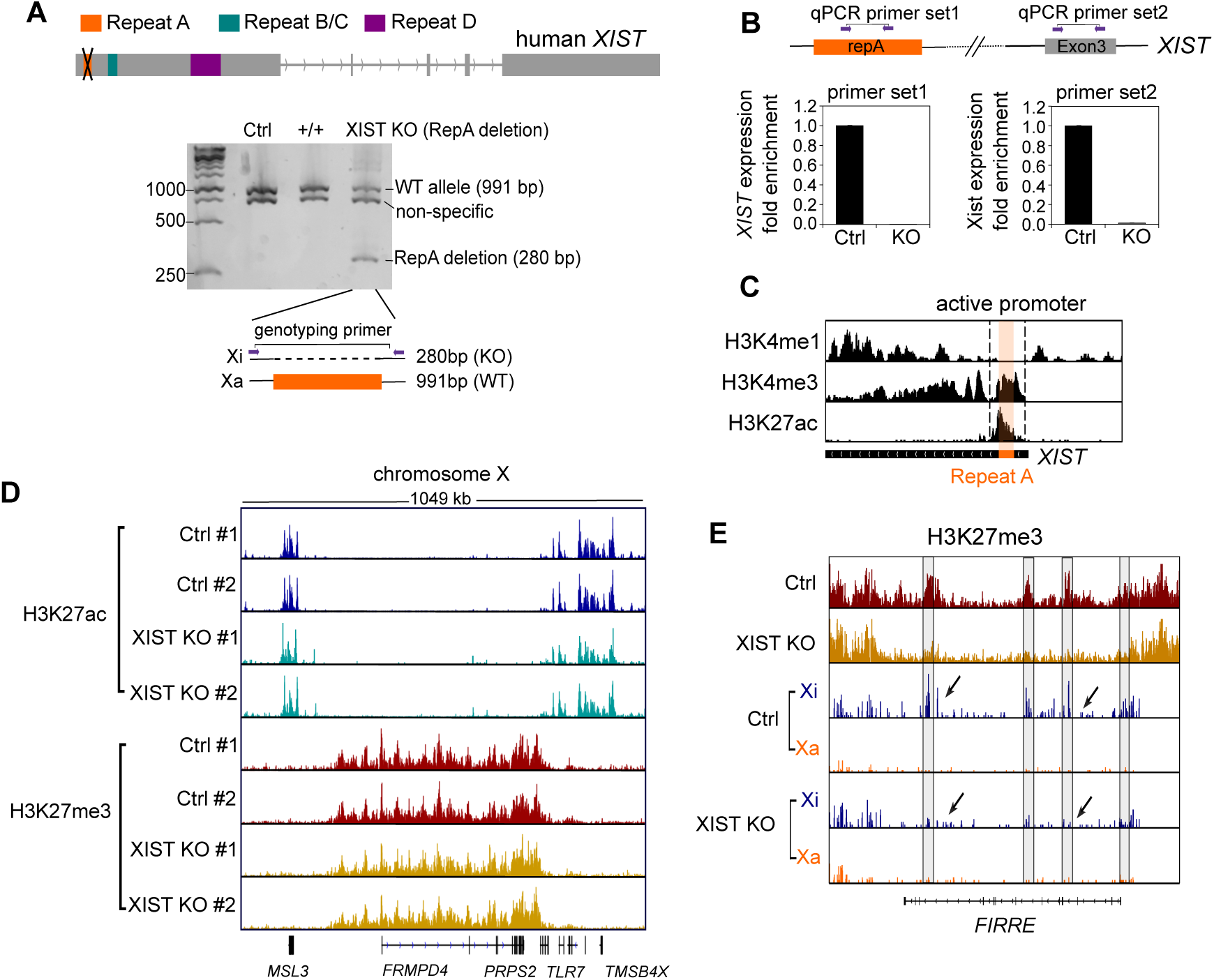
Allelic-specific ChIP-seq of H3K27ac and CUT&RUN of H3K27me3 in B cells with XIST repeat A deletion. Related to Figure 3. A. Schematic view of XIST repeat A deletion by Cas9-RNP. Genotyping PCR around XIST repeat A showing the deletion of XIST repeat A at one allele in a clonal XIST repeat A deletion cell line (XIST KO, third lane). B. qRT-PCR results showing the XIST expression in control and XIST KO cells using two primer sets targeting repeat A and exon3 respectively. C. Genome tracks of H3K4me1, H3K4me3 and H3K27ac of XIST gene in B cell line. Repeat A region is highlighted in orange. D. Genome tracks of replicates of H3K27ac ChIP-seq and H3K27me3 CUT&RUN in control and XIST KO group at chromosome X. E. H3K27me3 CUT & RUN track of XIST-dependent gene *FIRRE* between control and XIST KO group.

**Figure S4.**
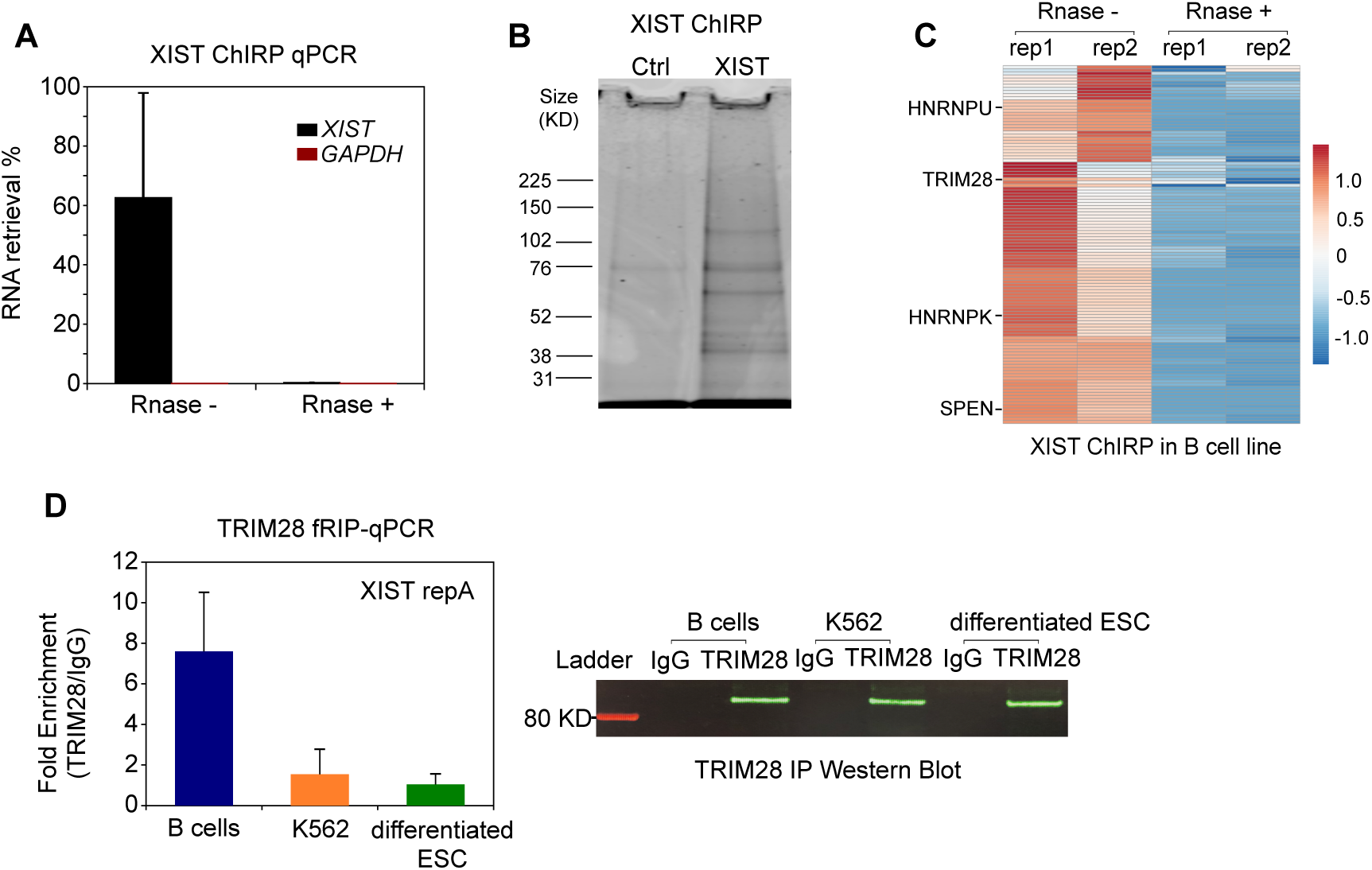
XIST ChIRP-MS specificity. Related to Figure 4. A. ChIRP-qPCR result showing the percentage of RNA retrieval of *XIST* and *GAPDH* in XIST pulled down portion compared to input RNA in B cell line. RNAse treatment prior to XIST pull-down is showed as a negative control. B. Coomassie blue stained gel showing the proteins pulled down by XIST probes and negative control with no XIST probes in B cell line. C. Heatmap showing the z score of peptide counts of XIST-associated proteins with or without Rnase treatment in B cell line. D. Bar plot showing the TRIM28 formaldehyde RIP-qPCR on A repeat of XIST in B cell line, myeloid K562 cell line, and differentiated ESCs. Western blot showing the pull down efficiency of TRIM28 in three cell types.

**Figure S5.**
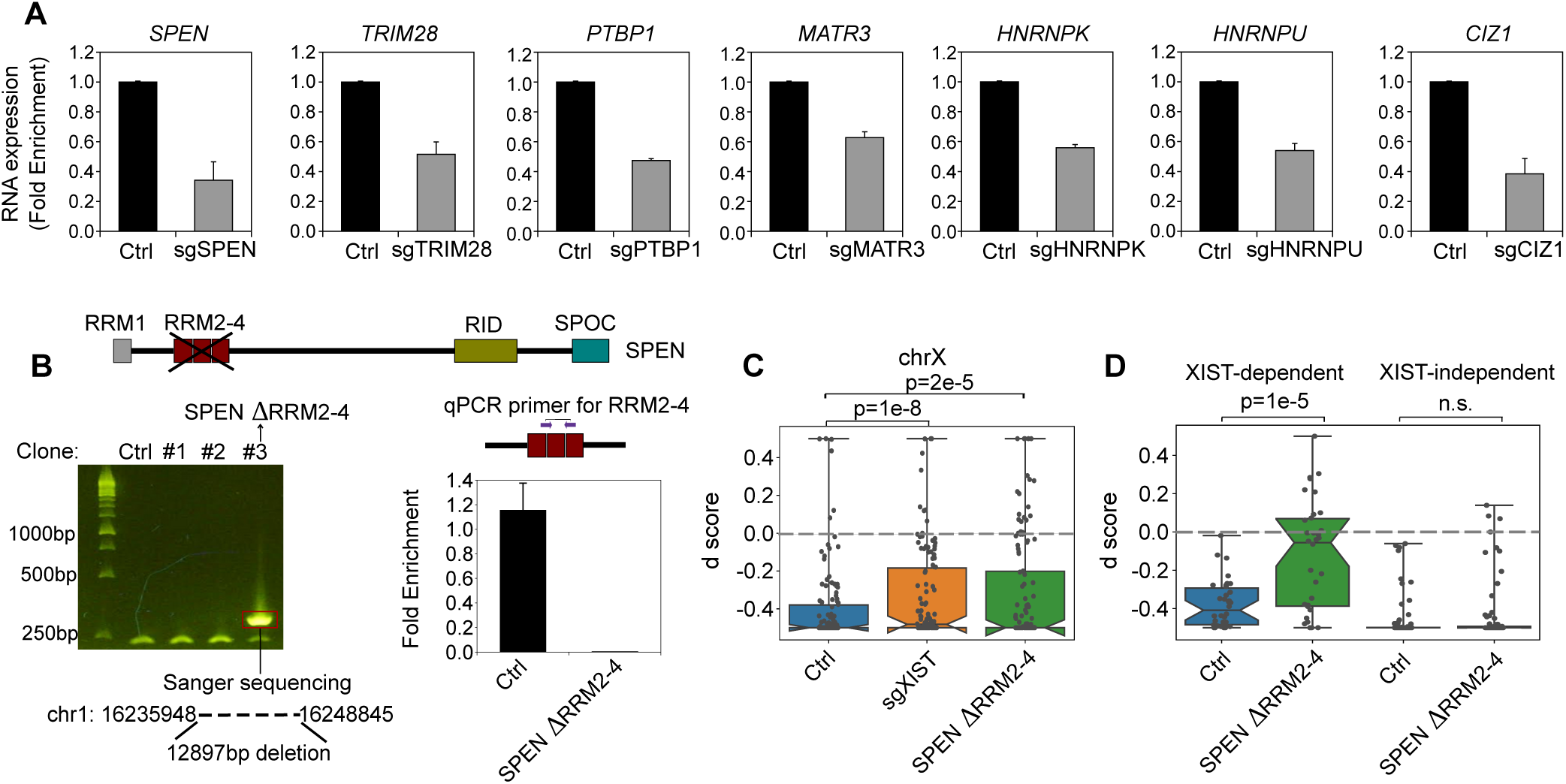
The perturbation efficiency of key XIST-cofactors. Related to Figure 5. A. qRT-PCR results showing the relative gene expression of specific XIST co-factors between non-targeting sgRNA perturbed cells (control) and XIST co-factors perturbed cells. Bar plots showing the mean value of RNA fold enrichment from two biological replicates. B. Genotyping PCR showing clones of B cell line with the deletion of SPEN RRM2-4 deletion (left). The qRT-PCR result showing the relative expression of SPEN RRM2-4 in control group and SPEN RRM2-4 deletion mutant group (right). C. Box plot showing the distribution of d score of allelic expression of genes on the chrX between control, sgXIST and SPEN mutant group. P value was calculated by paired t-test. D. Box plot showing the d score of allelic expression of XIST-dependent genes and XIST-independent genes in control and SPEN mutant group. P value was calculated by paired t-test.

**Figure S6.**
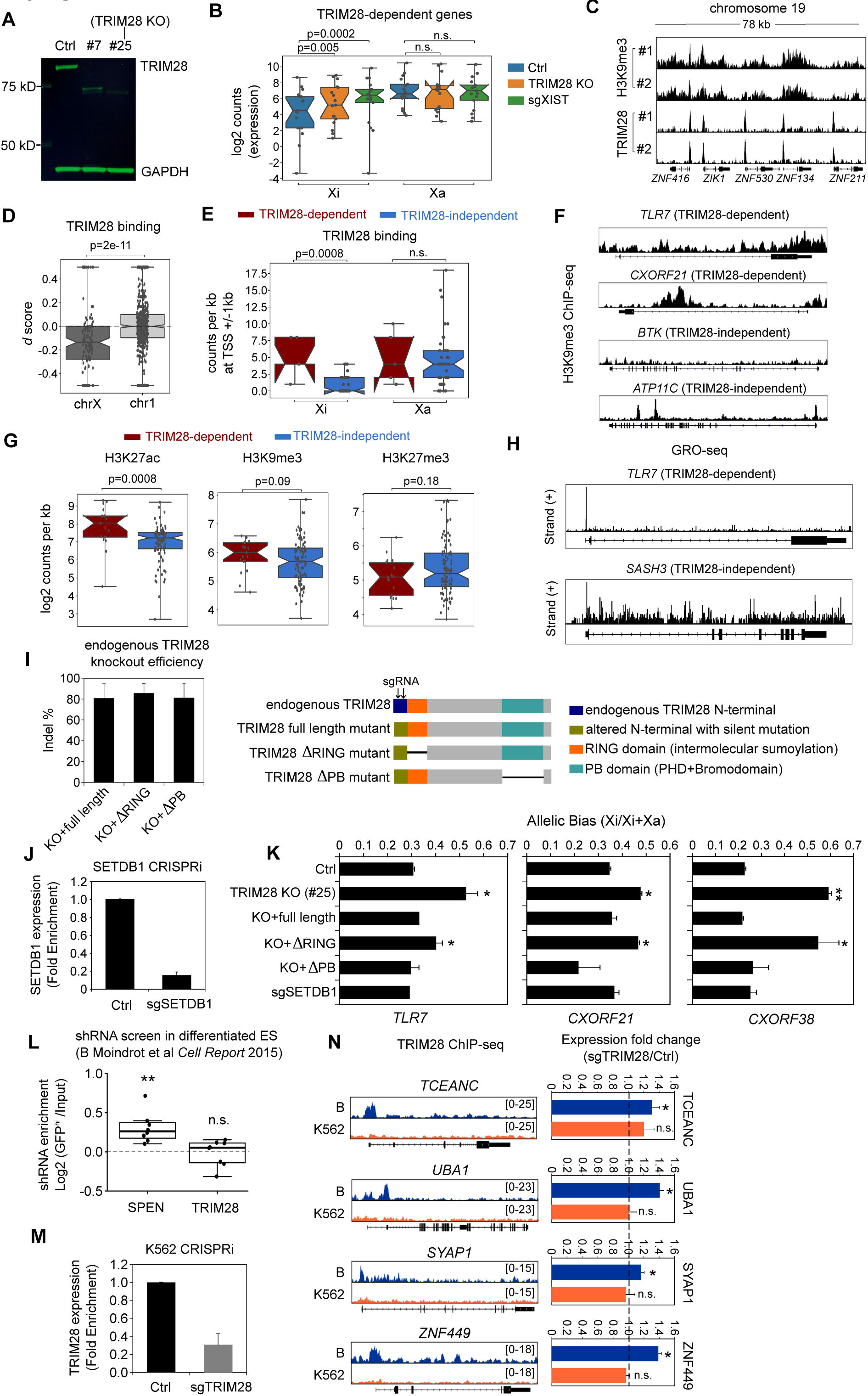
Allelic-specific RNA-seq in TRIM28 KO cells and TRIM28 ChIP-seq analysis. Related to Figure 6. A. Western blot showing the protein level of TRIM28 and GAPDH (internal control) in control and two clonal sgTRIM28 B cell lines (#7 and #25) by Cas9-RNP mediated knocking out. Over 99% TRIM28 proteins are depleted in #25 B cell line and this cell line was used as TRIM28 KO B cells. B. Box plot showing the log2 counts of expression of TRIM28-dependent genes at Xi and Xa in Ctrl, TRIM28 KO and sgXIST group. P value was calculated using paired t-test. C. Genome tracks showing the replicates of H3K9me3 CUT&RUN and TRIM28 ChIP-seq at ZNF gene clusters in B cell line. D. Box plot showing the allelic bias of TRIM28 binding at chrX and autosome chr1. P value was calculated via non-parametric Mann-Whitney test. E. Box plot showing the counts of TRIM28 binding at TSS +/- 1kb region of TRIM28- dependent (red) and TRIM28-independent genes (blue) between Xi and Xa. P value was calculated via non-parametric Mann-Whitney test. F. Genome tracks showing the H3K9me3 profiles of representative TRIM28-dependent and TRIM28-independent genes in B cell line. G. Box plots showing the occupancy density of H3K27ac, H3K9me3 and H3K27me3 at TSS +/- 1kb region of TRIM28-dependent (red) and TRIM28-independent genes (blue). P values were calculated via non-parametric Mann-Whitney test. H. Genome tracks showing the GRO-seq profile of nascent RNA at representative genes *TLR7* (TRIM28-dependent) and *SASH3* (TRIM28-independent) in B cell line. I. Bar plot showing the percentage of indels introduced after endogenous TRIM28 knockout via CRISPR-Cas9 editing in B cells that are overexpressed with TRIM28 full length, RING domain deletion, and PB domain deletion mutants (left). Schematic view of domains in endogenous TRIM28 and exogenous TRIM28 mutants. J. Bar plot showing the relative expression of SETDB1 in control and sgSETDB1 group using qRT-PCR. Mean value was calculated from two biological replicates. K. Bar plot showing the pyrosequencing result of three TRIM28-dependent genes (TLR7, CXORF21, CXORF38) in control B cells, TRIM28 KO cells, B cells that are first expressed with different TRIM28 mutants and then deplete endogenous TRIM28 with CRISPR-Cas9, B cells with SETDB1 perturbation. Mean value was calculated from two biological replicates. L. Box plot showing the SPEN and TRIM28 shRNA enrichment in GFP^hi^ cells compared to input cells using a published shRNA screen data in differentiated ES cells. M. Bar plot showing the relative expression of TRIM28 in K562 cells transduced with non-targeting control or sgTRIM28. Mean value was calculated from two biological replicates. N. Genome tracks showing TRIM28 ChIP-seq in GM B cells and K562 myeloid cells on 4 TRIM28-dependent genes (left). Bar plot showing the relative expression fold enrichment of TRIM28-dependent genes in sgTRIM28 group compared to control group in K562 cells (right). Mean value was calculated from two biological replicates. P value was calculated using unpaired t-test. * : p<0.05; **: p< 0.001.

**Figure S7.**
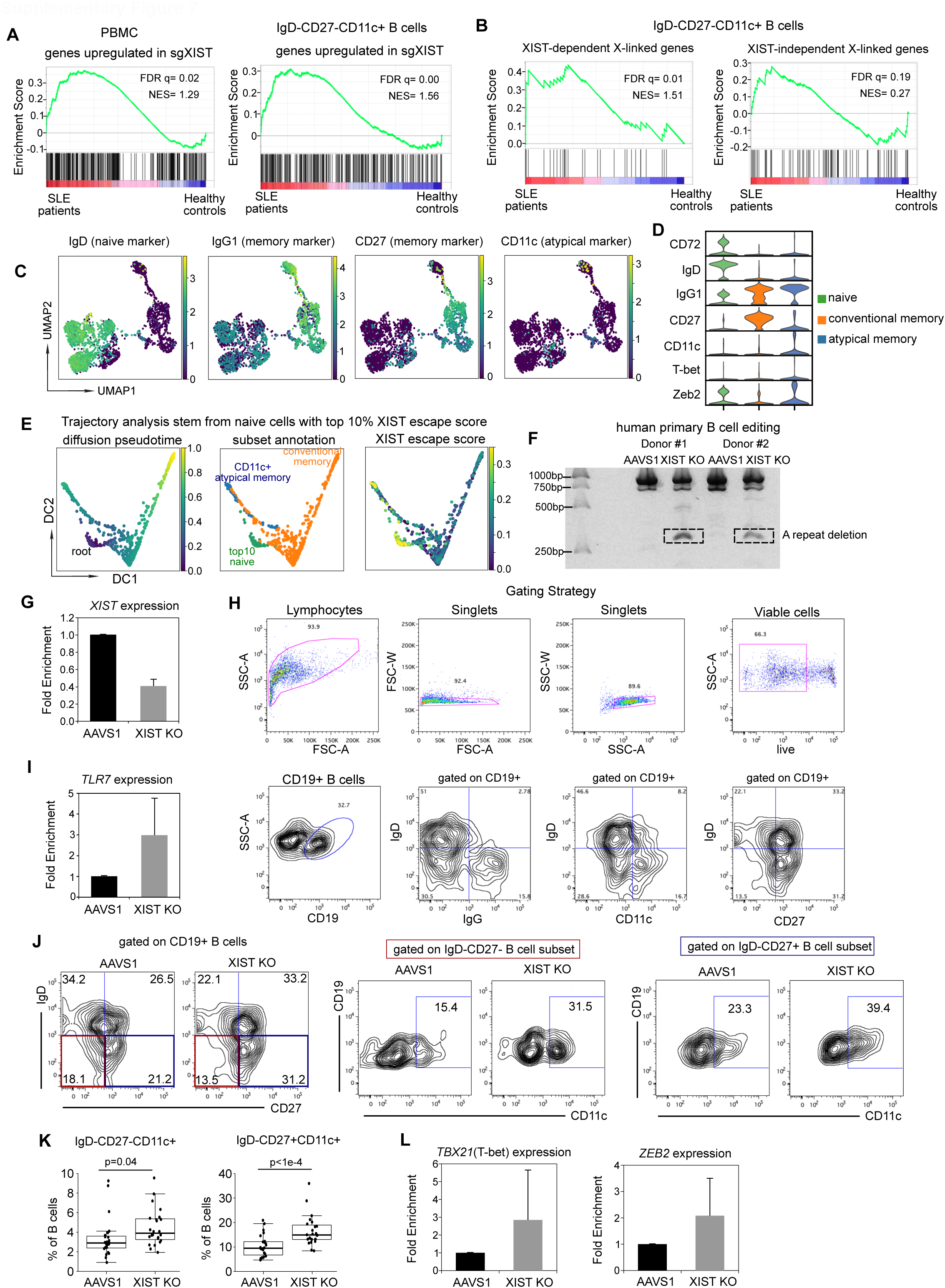
The Cas9-RNP mediated perturbation of XIST in human primary B cells. Related to Figure 7. A. GSEA analysis showing enrichment of gene set that is upregulated after XIST perturbation in PBMC or IgD-CD27-CD11c+ DN atypical B cells of female SLE patients compared to healthy donors. Normalized Enrichment Score (NES) and FDR q value are shown on the plot. B. GSEA analysis showing enrichment of XIST-dependent X-linked genes and XIST-independent X-linked genes in IgD-CD27-CD11c+ DN atypical B cells of female SLE patients compared to healthy donors. C. UMAP plots showing representative markers of naive (IgD), conventional memory (IgG1, CD27), and atypical memory (IgG1, CD11c) B cells. D. Stacked violin plots showing more markers for these three B cell subsets (naive B cell: CD72, IgD; conventional memory B cell: IgG1, CD27); atypical memory B cell: IgG1, CD11c, T-bet, Zeb2). E. Diffusion plots showing the pseudotime trajectory started from the naive cells with top10 percentile of XIST escape score. Left: diffusion pseudotime plot. Middle: subset annotated projected on diffusion map. Right: XIST escape score on diffusion map. F. Gel showing XIST A repeat deletion by genotyping PCR on Cas9-edited human primary B cells from two different female donors. The band highlighted on the gel showed A repeat deletion by sanger sequencing. G. qRT-PCR result showing *XIST* RNA expression after XIST repeat A deletion. H. Gating strategy of viable CD19+ B cells. I. qRT-PCR result showing *TLR7* RNA expression after XIST repeat A deletion. J. Flow cytometry analysis of IgD-CD27-CD11c+ and IgD-CD27+CD11c+ atypical memory B cells in AAVS1 and XIST KO group. K. Box plot showing the percentage of these B cell subsets among total B cells after XIST repeat A deletion. The data are pooled from 8 healthy donors with 2-3 replicates from each donor. P value was calculated by paired t-test. L. qRT-PCR result showing *Tbx21 and ZEB2* relative RNA expression after the loss of XIST.

## Notes

### Summary of Updates

Figure 1 and 7 are flattened due to formating issue

